# Bayesian model reveals latent atrophy factors with dissociable cognitive trajectories in Alzheimer’s disease

**DOI:** 10.1101/064295

**Authors:** Xiuming Zhang, Elizabeth C. Mormino, Nanbo Sun, Reisa A. Sperling, Mert R. Sabuncu, B.T. Thomas Yeo, for the Alzheimer’s Disease Neuroimaging Initiative

**Affiliations:** Department of Electrical and Computer Engineering, ASTAR-NUS Clinical Imaging Research Centre, Singapore Institute for Neurotechnology and Memory Networks Program, National University of Singapore, Singapore; Department of Neurology, Massachusetts General Hospital/Harvard Medical School, Charlestown, MA 02129, USA; Martinos Center for Biomedical Imaging, Massachusetts General Hospital/Harvard Medical School, Charlestown, MA 02129, USA; Computer Science and Artificial Intelligence Laboratory, Massachusetts Institute of Technology, Cambridge, MA 02138, USA; Centre for Cognitive Neuroscience, Duke-NUS Graduate Medical School, Singapore

**Author notes:** Address correspondence to: B.T. Thomas Yeo ECE, ASTAR-NUS CIRC, SINAPSE & MNP National University of Singapore. Data used in preparation of this article were obtained from the Alzheimer’s Disease Neuroimaging Initiative (ADNI) database (http://adni.loni.usc.edu). As such, the investigators within the ADNI contributed to the design and implementation of ADNI and/or provided data but did not participate in analysis or writing of this report. A complete listing of ADNI investigators can be found in **Supporting Information.**.

## Abstract

We employed a data-driven Bayesian model to automatically identify distinct latent factors of overlapping atrophy patterns from voxelwise structural magnetic resonance imaging (MRI) of late-onset Alzheimer’s disease (AD) dementia patients. Our approach estimated the extent to which multiple distinct atrophy patterns were expressed within each participant rather than assuming that each participant expressed a single atrophy factor. The model revealed a temporal atrophy factor (medial temporal cortex, hippocampus and amygdala), a subcortical atrophy factor (striatum, thalamus and cerebellum), and a cortical atrophy factor (frontal, parietal, lateral temporal and lateral occipital cortices). To explore the influence of each factor in early AD, atrophy factor compositions were inferred in beta-amyloid-positive (Aβ+) mild cognitively impaired (MCI) and cognitively normal (CN) participants. All three factors were associated with memory decline across the entire clinical spectrum, whereas the cortical factor was associated with executive function decline in Aβ+ MCI participants and AD dementia patients. Direct comparison between factors revealed that the temporal factor showed the strongest association with memory, while the cortical factor showed the strongest association with executive function. The subcortical factor was associated with the slowest decline for both memory and executive function compared to temporal and cortical factors. These results suggest that distinct patterns of atrophy influence decline across different cognitive domains. Quantification of this heterogeneity may enable the computation of individual-level predictions relevant for disease monitoring and customized therapies. Code from this manuscript is publicly available at link_to_be_added.

## Significance

Alzheimer’s disease (AD) is the most common form of dementia. Heterogeneity in AD complicates efforts for diagnosis and disease monitoring. Here we model the heterogeneity of atrophy in AD patients, demonstrating for the first time that most AD patients and at-risk nondemented participants express multiple latent atrophy factors to varying degrees. Furthermore, these atrophy factors are associated with distinct decline trajectories of memory and executive function, highlighting the relevance of these atrophy patterns in understanding the clinical course of AD dementia. Our results provide a framework by which biomarker readouts could predict disease progression at the individual level. Our analytic strategy could potentially be utilized to discover subtypes within and across other heterogeneous brain disorders, such as autism and schizophrenia.

## Introduction

Alzheimer’s disease (AD) dementia is a devastating neurodegenerative disease that affects 11% of individuals over age 65 with no disease modifying treatment available. Accurate *in vivo* biomarkers are urgently needed to assist in early detection of at-risk individuals, improve diagnosis, monitor disease progression, and serve as outcome measures in clinical trials.

Although AD is typically associated with an amnestic clinical presentation and disruption of the medial temporal lobe (Salmon & Bondi, 2009), it has become increasingly clear that heterogeneity exists within this disease. Specifically, heterogeneity has been observed in the clinical presentation of AD (Lam et al., 2013), the spatial distribution of neurofibrillary tangles (NFT) (Murray et al., 2011, Ossenkoppele et al., 2016), as well as the presence of co-morbid pathologies such as vascular disease, Lewy bodies and TDP-43 (Schneider et al., 2009, Josephs et al., 2014). Interestingly, the spatial distribution of atrophy varies across AD subtypes defined on the basis of NFT distribution (Whitwell et al., 2012), suggesting that analyses of gray matter patterns are useful to characterize heterogeneity in AD. Furthermore, although distinct atrophy patterns have been observed in patients that clearly show atypical clinical presentations (Ossenkoppele et al., 2015), heterogeneity in gray matter atrophy has also been reported among late-onset AD cases (Dickerson & Wolk, 2011). It is therefore likely that the ability to quantify varying patterns of atrophy among AD patients will help inform our understanding of fundamental disease processes.

In this study, we sought to explore the heterogeneity of atrophy patterns in late-onset AD using a data-driven Bayesian framework that accounted for and estimated *latent* AD atrophy factors derived from structural MRI data. The mathematical framework that we employed, latent Dirichlet allocation (LDA; Blei et al., 2003), has been successfully utilized to extract overlapping brain networks from functional MRI (Yeo et al., 2014) and meta-analytic data (Yeo et al., 2015; Bertolero et al., 2015). Importantly, this approach does not require the atrophy pattern of an individual to be determined by a single atrophy factor. Instead, the model allows the possibility that multiple latent factors are expressed to varying degrees within an individual. For example, the atrophy pattern of a patient might be 90% due to factor 1 and 10% factor 2, while the atrophy pattern of another patient might be 60% due to factor 1 and 40% factor 2. Given that multiple contributors that are not mutually exclusive may influence heterogeneity in AD, such as the spatial location of NFT pathology (Murray et al., 2011, Ossenkoppele et al., 2016), coexisting non-AD pathologies (Schneider et al., 2009) and genetics (Dickerson & Wolk, 2011), we believe it is more biologically plausible that individuals express varying degrees of distinct atrophy factors rather than one single factor. Thus, the LDA approach is particularly well suited for these analyses and will provide insight into whether expressing multiple atrophy factors is common among late-onset AD patients.

Most studies investigating the heterogeneity of AD have examined patients soon after AD onset or at advanced AD stages (e.g., Murray et al., 2011, Whitwell et al., 2012, Noh et al., 2014, Byun et al., 2015, Scheltens et al., 2015). However, the pathophysiological processes of AD begin at least a decade before clinical diagnosis (Villemagne et al., 2013), suggesting that the emergence of this heterogeneity may occur prior to the onset of clinical dementia. In this study, we therefore examined how distinct atrophy factors identified in AD dementia patients were associated with longitudinal cognitive decline early in nondemented participants that were at risk for AD dementia based on elevated beta-amyloid (Sperling et al., 2011, Albert et al., 2011, Rowe et al., 2013).

Our study makes three significant contributions. First, we introduced an innovative modeling strategy where expressions of multiple atrophy patterns are estimated rather than assigning each participant to a single subtype. Second, our approach harnesses the rich multidimensional information across all gray matter voxels, avoiding the need for *a priori* selection of regions and enabling an in-depth exploration of atrophy patterns. Finally, application of this approach to participants spanning the clinical spectrum revealed that latent atrophy factors are associated with distinct memory and executive function trajectories, providing novel insights into the impact of disease heterogeneity throughout the prolonged course of AD.

## Results

### Overall Approach

Our approach involved three main steps. In **Step I**, we performed latent Dirichlet allocation (a Bayesian model; Blei et al., 2003) to estimate latent atrophy factors in 188 AD dementia patients and used this model to extract factor compositions in two independent samples of nondemented participants: 147 beta-amyloid-positive (Aβ+) mild cognitively impaired (MCI) and 43 Aβ+ cognitively normal (CN) participants. In **Step II**, we examined robustness across different analytic approaches and investigated characteristics of the factor compositions across participants. Finally, in **Step III** we examined the associations between atrophy factors and different cognitive domains (memory and executive function). The results of each step are described in detail below.

#### I. Discovering Latent Atrophy Factors in AD Dementia Patients

We employed the Bayesian latent Dirichlet allocation (LDA) model (Blei et al., 2003) to encode our assumption that a patient expresses one or more *latent* atrophy factors (Fig. 1). The LDA model was applied to the structural MRI of 188 AD dementia patients. Given the voxelwise gray matter density values derived from structural MRI (FSL-VBM; Douaud et al., 2007) and a predefined number of factors K, the model is able to estimate the probability that a particular factor is associated with atrophy at a specific spatial location (i.e., Pr(Voxel | Factor) or probabilistic atrophy map of the factor) and the probability that an individual expresses each atrophy factor (i.e., Pr(Factor | Patient) or atrophy factor composition of the individual). Importantly, resulting atrophy factors were not predetermined, but estimated from data (see **Methods**).

**Fig. 1.**
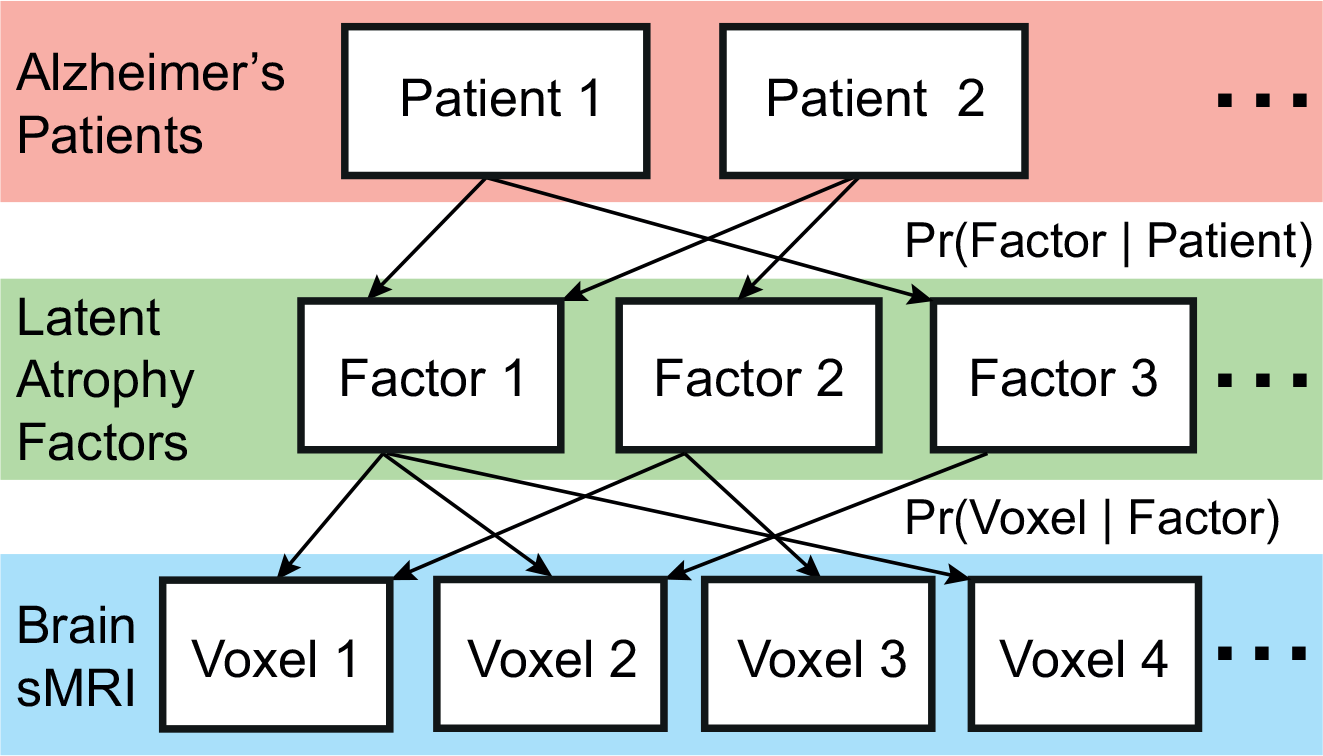
A Bayesian model of AD dementia patients, latent atrophy factors, and brain structural MRI. Underpinning our approach is the premise that each participant expresses one or more latent factors. Each factor is associated with distinct but possibly overlapping patterns of brain atrophy. The framework can be instantiated with a mathematical model (latent Dirichlet allocation; Blei et al., 2003), whose parameters can be estimated from the structural MRI data of AD dementia patients. The estimated parameters are the probability that a patient expresses a particular factor, i.e., Pr(Factor |Patient), and the probability that a factor is associated with atrophy at a MRI voxel, i.e., Pr(Voxel | Factor).

An important model parameter is the number of latent atrophy factors K.
Therefore, we first determined how factor estimation changed from K = 2 to 10. Visual inspection of the spatial distribution of each atrophy factor suggested that factor estimates from K = 2 through 10 were organized in a hierarchical fashion (Figs. 2 and S1). For instance, the two-factor model revealed one factor associated with atrophy in temporal and subcortical regions (“temporal+subcortical” Fig. 2A1) and another factor associated with atrophy throughout cortex (“cortical” Fig. 2A2). The three-factor model resulted in a similar cortical factor (Fig. 2B3 and Table S1C), while the temporal+subcortical factor split into a “temporal” factor associated with extensive atrophy in the medial temporal lobe (Fig. 2B1 and Table S1A) and a “subcortical” factor associated with atrophy in the cerebellum, striatum and thalamus (Fig. 2B2 and Table S1B). Likewise, the four-factor model resulted in the cortical factor splitting into “frontal cortical” and “posterior cortical” factors, whereas the temporal and subcortical factors remained the same (Fig. 2C). Sagittal and axial slices of these probabilistic atrophy maps are available in Fig. S1 in the **Supporting Information**(**SI**).

**Fig. 2.**
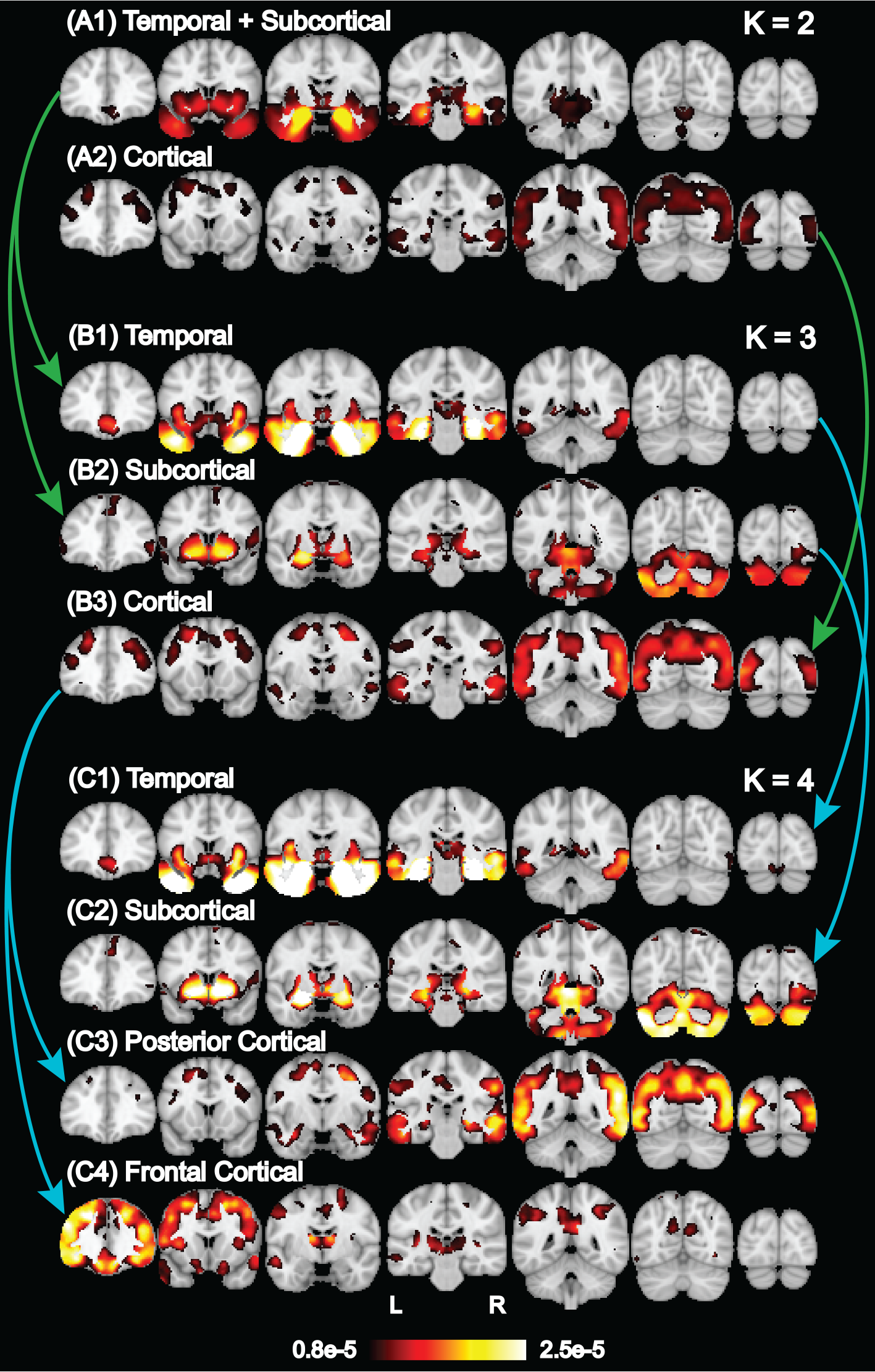
Hierarchy of latent atrophy factors with distinct atrophy patterns in AD. Bright color indicates higher probability of atrophy at that spatial location for a particular atrophy factor, i.e., Pr(Voxel | Factor). Each of the (A) two, (B) three and (C) four factors was associated with a distinct pattern of brain atrophy and was named accordingly. A nested hierarchy of atrophy factors was observed even though the model did not mandate such a hierarchy. For example, when going from two to three factors, the temporal+subcortical factor (A1) split into temporal (B1) and subcortical (B2) factors, while the cortical factor remained the same (A2 and B3). This hierarchical phenomenon was quantified for two to ten factors (Fig. S2).

To quantify the hierarchical phenomenon, we employed an exhaustive search to assess the possibility that two unknown factors in the (K+1)-factor model were subdivisions of an unknown factor in the K-factor model (while the other factors remained the same). The exhaustive search yielded a hypothesized factor hierarchy with associated correlation values quantifying the subdivision quality (see **Supplemental Methods** of **SI**). The high correlation values (Fig. S2) confirmed that additional factors emerged as subdivisions of lower-order factors, corresponding to a nested hierarchy of atrophy factors.

This nested hierarchy suggested that specification of different numbers of estimated factors might yield distinct insights into AD. In the remainder of this paper, we highlighted the results of three-factor model (Fig. 2B), since the emergence of the temporal and cortical factors were consistent with the “limbic-predominant” and “hippocampal-sparing” pathologically-defined AD subtypes previously reported (Murray et al., 2011, Whitwell et al., 2012). We additionally repeated analyses for two-and four-factor models, which yielded behavioral insights consistent with the three-factor model. These additional results are reported in **Supplemental Figures** of **SI**.

To explore the influence of atrophy factors in early AD, probabilistic atrophy maps Pr(Voxel | Factor) estimated from the AD dementia patients were used to infer factor compositions Pr(Factor | Participant) of the 190 Aβ+ nondemented participants using the standard variational expectation-maximization algorithm (Blei et al., 2003).

#### II. Examining Factor Robustness and Characteristics of Factor Compositions

Among the 188 AD dementia patients, 100 had their cerebrospinal fluid (CSF) amyloid data available. 91 of the 100 patients were Aβ+ (CSF amyloid concentration < 192 pg/ml; Shaw et al., 2009). We performed LDA on the subset of Aβ+ AD dementia patients (and on Aβ+ MCI participants; see **Supplemental Results** of **SI**) and compared atrophy patterns of the resulting factors with those derived using the larger sample (Fig. S3). Atrophy factors were similar across these methods, with an average correlation across all pairwise comparisons of r = 0.89. Given this similarity and to improve our estimates of the atrophy factors, we elected to use the atrophy factors derived from the larger sample of 188 AD dementia patients for subsequent analyses. Furthermore, resulting atrophy patterns were consistent between FreeSurfer (Fischl, 2012) and FSL-VBM, suggesting that the atrophy factors were robust to variations in image preprocessing software (see **Supplemental Results** and **Supplemental Methods** of **SI**).

To determine whether expression of atrophy factors remained stable over time, we examined the subset of ADNI 1 participants that had a two-year follow-up scan available (N = 560 out of 810). We were specifically interested in whether atrophy factors reflected different disease stages rather than different atrophy subtypes (for instance, high expression of the temporal factor may lessen over time with greater expression of the cortical factor). Therefore, we compared factor compositions after two years with baseline compositions. The factor probabilities were positivity correlated and highly consistent (r > 0.85 across all three factors, Fig. 3; see Fig. S4 for results by diagnostic group with additional amyloid information), suggesting that these factors do not merely reflect a sequence of atrophy patterns.

**Fig. 3.**
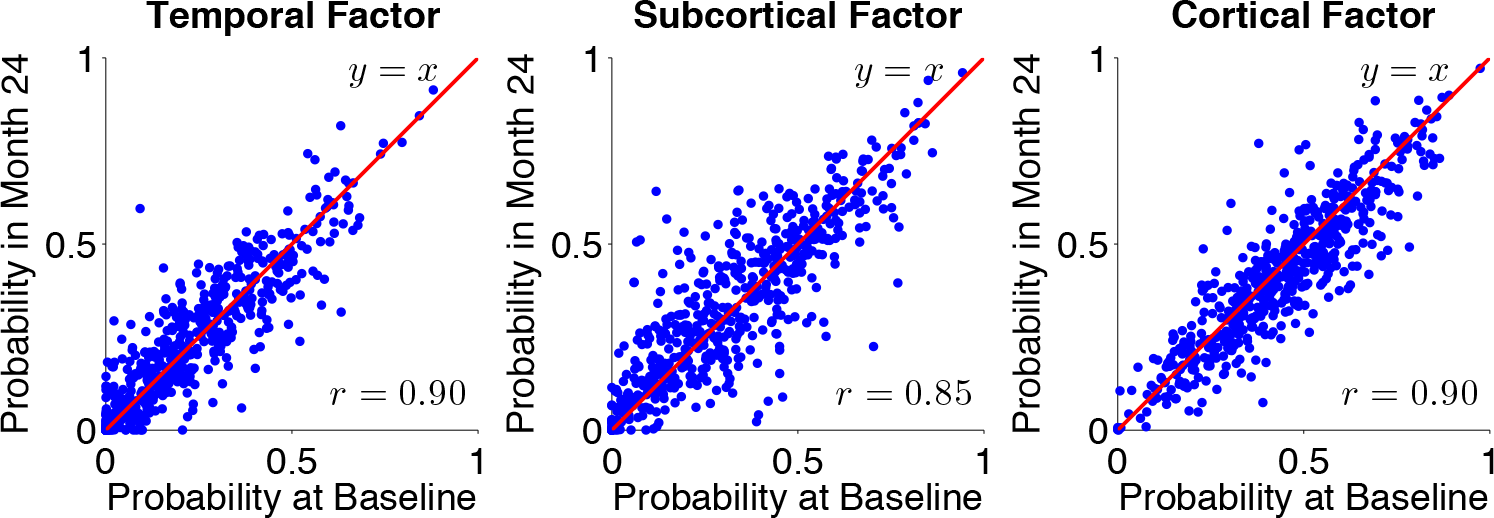
Stability of factor compositions over two years. Each blue dot represents a participant. For each plot, x-axis and y-axis represent, respectively, the probabilities of factor at baseline and two years after baseline. In the ideal case where factor probability estimates remain exactly the same after two years, one would expect a y = x linear fit as well as a r = 1 correlation. In our case, the linear fits were close to y = x with r > 0.85 for all three atrophy factors, suggesting that the factor compositions were stable despite disease progression.

Examination of atrophy factor compositions among AD dementia patients revealed that the majority expressed multiple latent atrophy factors rather than predominantly expressing a single atrophy factor (Fig. 4). Examination of factor compositions of the 190 Aβ+ nondemented participants revealed a similar pattern, such that the majority of participants expressed multiple atrophy factors (Fig. S5A). Factor compositions for the two-and four-factor models also suggest that most participants expressed multiple atrophy factors (Figs. S5B and S5C).

**Fig. 4.**
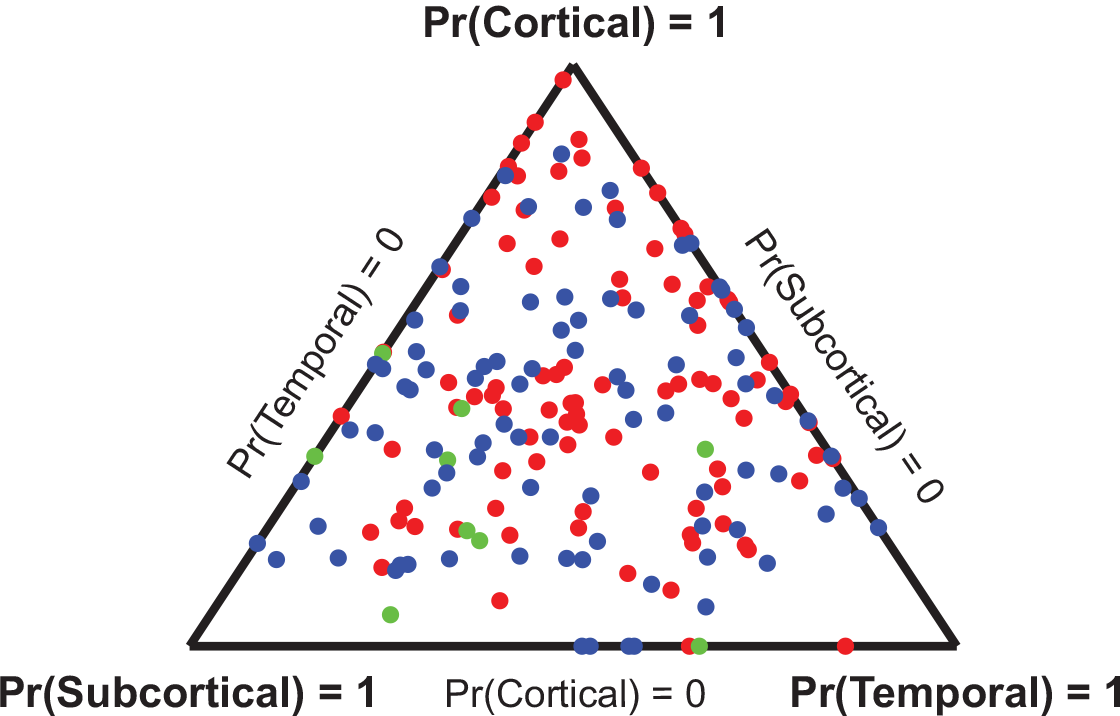
Factor compositions of 188 AD dementia patients. Each patient corresponds to a dot, whose location (in barycentric coordinates) represents the factor composition. Color indicates amyloid status: red for Aβ+, green for Aβ−, and blue for unknown. Corners of the triangle represent “pure factors”; closer distance to the respective corner indicates higher probability for the respective factor. Most dots are far from the corners, suggesting that most patients expressed multiple factors.

To understand the association between atrophy factors and demographic variables, general linear model (GLM; for continuous variables) and logistic regression (for binary variables) were conducted in the 188 AD dementia patients (Table S2). Briefly, the response variable was the variable of interest (e.g., age at AD onset), and the explanatory variables consisted of two columns encoding participants’ loading on the cortical and subcortical factors. The temporal factor was implicitly modeled because the factor probabilities summed to 1 (see **Methods**).

There were no significant differences in years from AD onset, education, sex or APOE ε4 loadings across the three factors. Importantly, amyloid level was not significantly different across factors. The cortical factor was associated with significantly younger baseline age than the temporal factor (p = 1e-5) and subcortical factor (p = 2e-6) as well as younger age at AD onset than the temporal factor (p = 3e-4) and subcortical factor (p = 7e-6). In addition, the subcortical factor was associated with higher APOE ε2 loading than the temporal factor (p = 0.01) and cortical factor (p = 0.04), but these were not significant when corrected for multiple comparisons.

Similar analyses were conducted for the Aβ+ MCI and CN groups. The only significant association was that among Aβ+ MCI participants, the cortical factor was associated with younger age at baseline compared to the temporal factor (p = 0.05) and subcortical factor (p = 0.02). However, this association did not survive after correcting for multiple comparisons.

#### III. Examining Associations Between Atrophy Factors and Cognition

We first examined diagnostic group differences in memory (ADNI-Mem; Crane et al., 2012) and executive function (ADNI-EF; Gibbons et al., 2012) without considering factor compositions. As expected, cross-sectional memory was worse for AD dementia patients (mean = −0.84) compared to Aβ+ MCI participants (mean = −0.21; t-test p = 5e-23). Aβ+ MCI participants had worse memory than Aβ+ CN participants (mean = 0.93; t-test p = 2e-26). Likewise, cross-sectional executive function was worse for AD dementia patients (mean = −0.92) compared to Aβ+ MCI participants (mean = −0.17; t-test p = 3e-16). Aβ+ MCI participants had worse executive function than Aβ+ CN participants (mean = 0.50; t-test p = 1e-7).

We then examined a GLM predicting *cross-sectional* memory and executive function, which included both diagnosis and factor compositions as well as their interactions as predictors (Figs. 5A1 and 5B1; see **Methods** for model details). This analysis revealed that all factors were associated with baseline memory, and these associations continued to worsen across the disease spectrum (Fig. 5A1). For crosssectional executive function, there was only an association with the cortical factor, and this association also worsened across the disease spectrum (Fig. 5B1).

Next we examined a linear mixed-effects (LME) model predicting *longitudinal* change in memory and executive function (Figs. 5A2 and 5B2). The LME model provides significantly improved exploitation of longitudinal measurements (Bernal-Rusiel et al., 2012) by accounting for both intra-individual measurement correlations and inter-individual variability. The model setup was the same as the GLM above, except that time and its interactions with diagnosis and factor compositions were included as predictors (see **Supplemental Methods** of **SI**).

This analysis revealed that the temporal and subcortical factors exhibited memory decline that began in CN and maintained similar memory decline rates in MCI and AD (Fig. 5A2). In contrast, the cortical factor was not associated with memory decline in CN, but demonstrated faster decline in MCI compared to CN and in AD compared to MCI (Fig. 5A2). The cortical factor was not associated with executive function decline in CN, but showed faster longitudinal executive function decline in MCI compared to CN and in AD compared to MCI (Fig. 5B2).

**Fig. 5.**
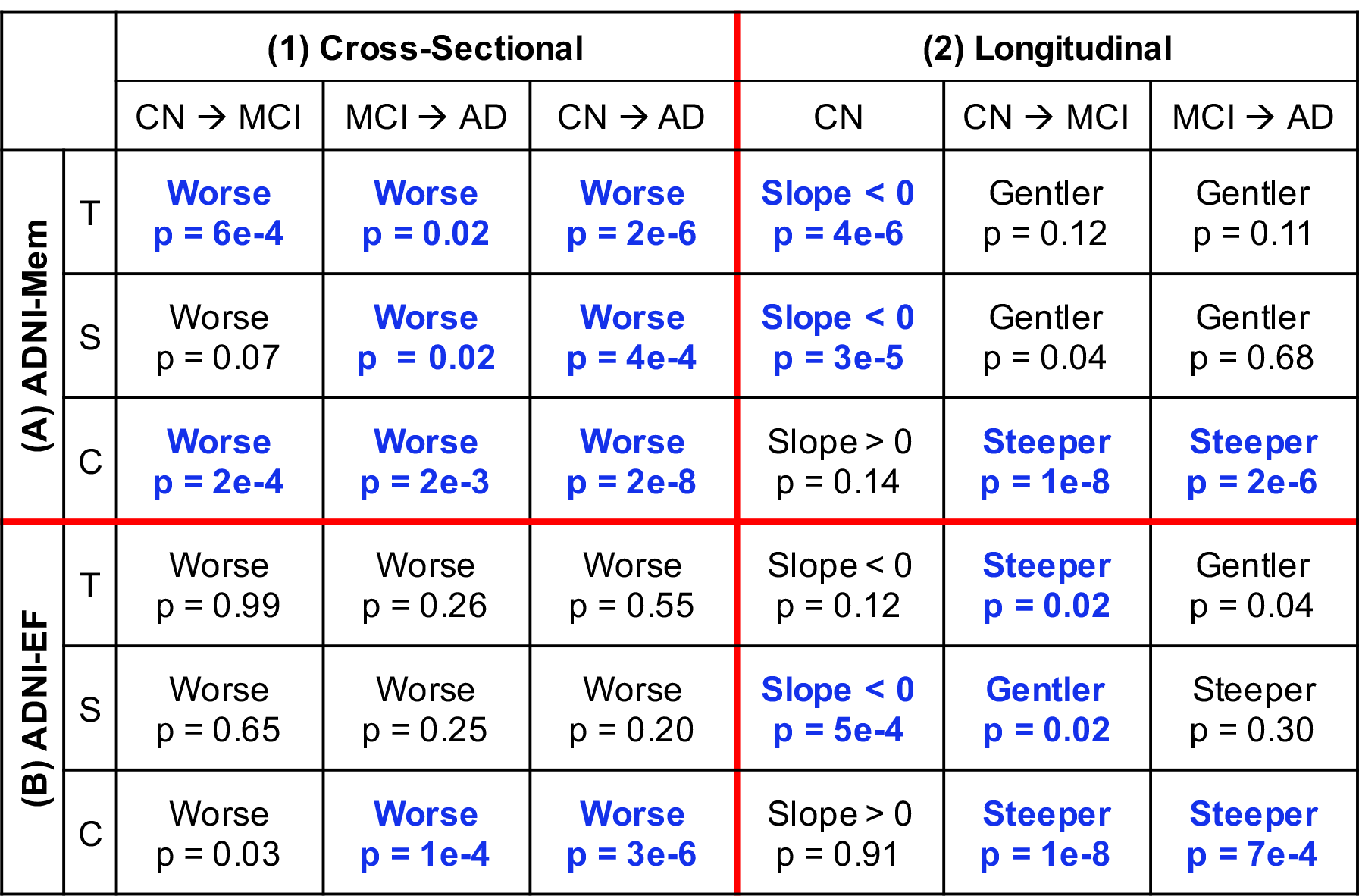
Differences by diagnosis and atrophy factor in (1) cross-sectional baseline and (2) longitudinal decline rates of (A) memory and (B) executive function. Comparisons remaining significant after false discovery rate (FDR; q = 0.05) control are highlighted in blue. “T”, “S” and “C” are short for temporal, subcortical and cortical factors, respectively. For example, the top left cell of (A1) suggests that Aβ+ MCI participants with high loading on the temporal factor had worse baseline memory than Aβ+ CN participants with high loading on the same factor (p = 6e-4). On the other hand, the bottom left cell of (B2) suggests that Aβ+ CN participants expressing the cortical factor did not exhibit executive function decline (p = 0. 91), while the bottom right cell of (B2) suggests that AD dementia patients expressing the cortical factor showed faster executive function decline than Aβ+ MCI participants expressing the same factor (p = 7e-4).

In our final set of analyses examining cognition, we directly compared the three factors. The GLM and LME models were exactly the same as the previous sections, but we instead focused on the contrasts between factors.

For cross-sectional memory, the temporal factor was associated with worse performance than the subcortical (p=3e-6) and cortical (p = 7e-3) factors among AD dementia patients (Fig. 6A). Similar results were found for Aβ+ MCI participants (Fig. S7A1). Among Aβ+ CN participants, there was no memory difference across the atrophy factors (Fig. S7A1). For cross-sectional executive function, the cortical factor was associated with worse performance than the temporal (p = 0.01) and subcortical (p = le-5) factors among AD dementia patients (Fig. 6B). There was no executive function difference across the factors among Aβ+ CN and MCI participants (Fig. S7B1).

**Fig. 6.**
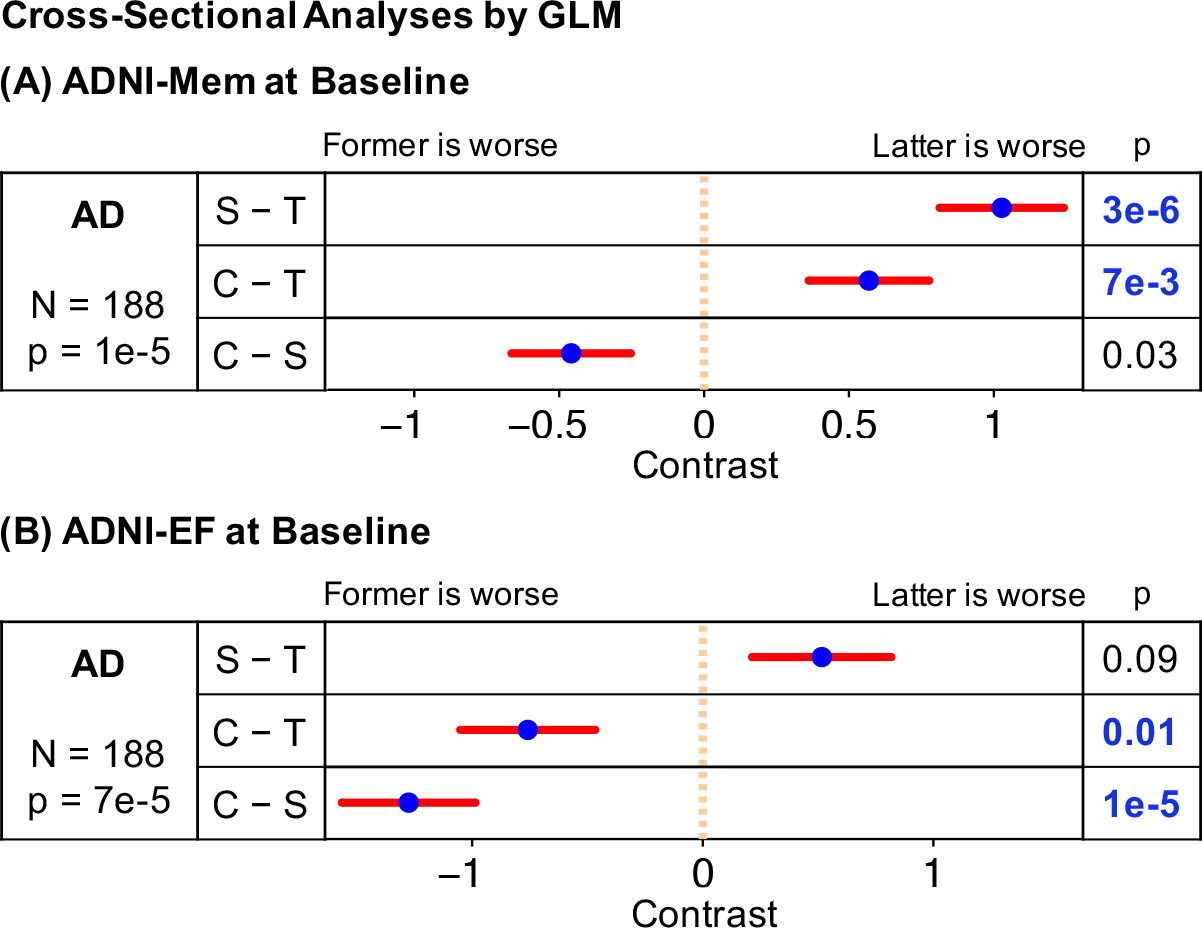
Comparisons of baseline (A) memory and (B) executive function in AD dementia patients across factors. Comparisons remaining significant after FDR (q = 0.05) control are highlighted in blue. “T”, “S” and “C” are short for the temporal, subcortical and cortical factors, respectively. Blue dots are estimated differences between “pure atrophy factors”, and red bars show the standard errors (see **Methods**). For example, the top row in (A) suggests that AD dementia patients expressing the temporal factor had worse baseline memory than AD dementia patients expressing the subcortical factor (p = 3e-6).

For longitudinal change in memory (Fig. 7A), the cortical factor was associated with faster longitudinal memory decline than the temporal (p = le-4) and subcortical (p = 4e-6) factors among AD dementia patients. Among Aβ+ MCI participants, the subcortical factor was associated with slower decline rate than the cortical (p = 8e-4) and temporal (p = 4e-3) factors. Finally, among Aβ+ CN participants, the cortical factor showed slower memory decline than the temporal (p = le-4) and subcortical (p = 3e-3) factors.

For longitudinal change in executive function (Fig. 7B), the cortical factor was associated with faster executive function decline than temporal (p = 2e-3) and subcortical (p = 2e-4) factors among AD dementia patients. Among Aβ+ MCI participants, the subcortical factor had slower decline than the cortical (p = 8e-9) and temporal (p = le-8) factors. There was no executive function decline difference across the factors among Aβ+ CN participants.

**Fig. 7.**
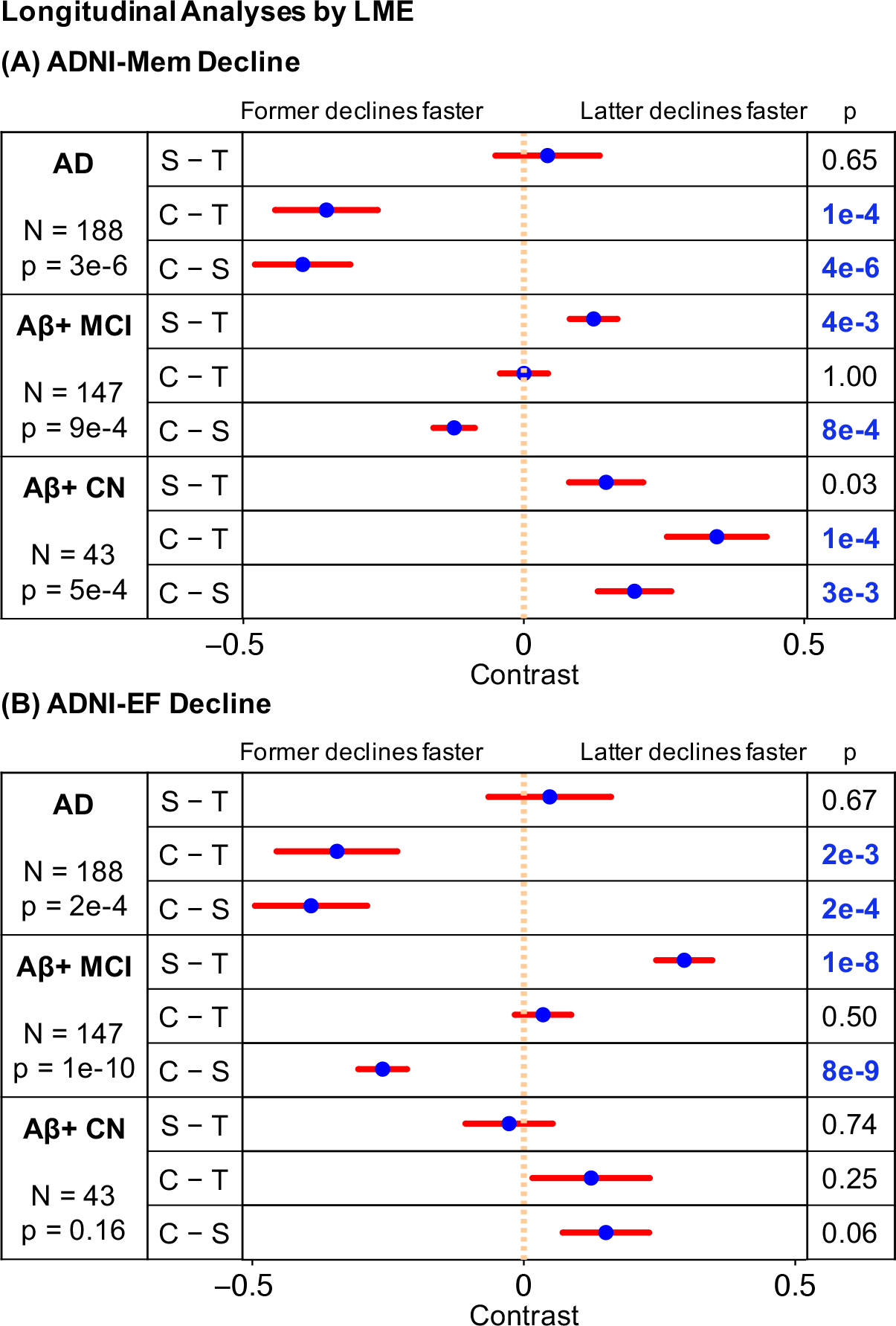
Comparisons of (A) memory and (B) executive function decline rates across factors by clinical group. Comparisons remaining significant after FDR (q = 0.05) control are highlighted in blue. “T”, “S” and “C” are short for the temporal, subcortical and cortical factors, respectively. Blue dots are estimated differences between “pure atrophy factors”, and red bars show the standard errors (see **Supplemental Methods** of **SI**). For example, the second row in (A) suggests that AD dementia patients expressing the cortical factor showed faster memory decline than patients expressing the temporal factor (p = 1e-4).

All cognitive analyses were repeated using the two-and four-factor LDA atrophy factors (Figs. S6 and S8). The results were consistent with the three-factor model (**Supplemental Results** of **SI**). In addition, associations between mini-mental state examination (MMSE) and the three atrophy factors are reported in Fig. S7C.

**Fig. 8.**
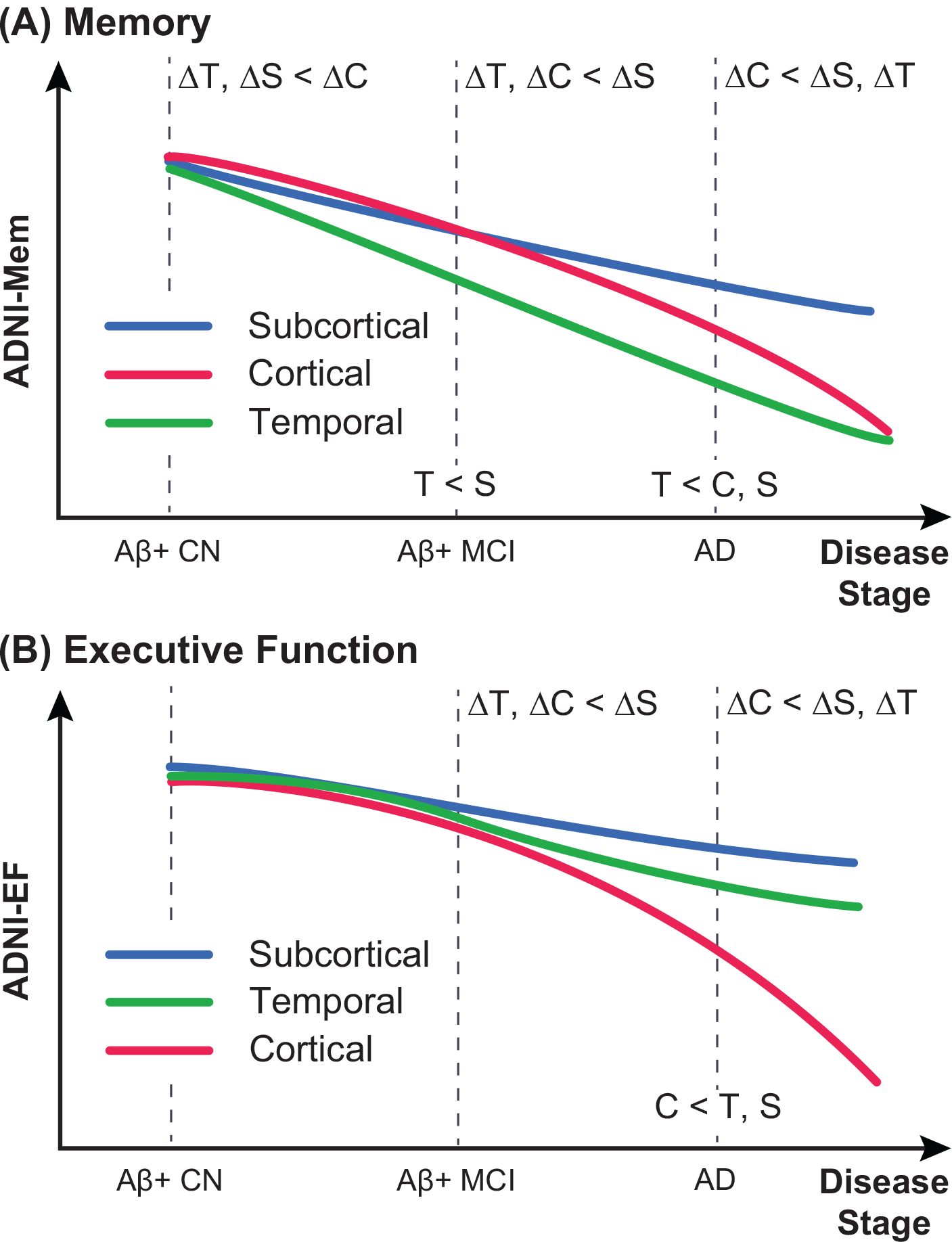
Schematics of distinct (A) memory and (B) executive function trajectories for temporal, subcortical and cortical atrophy factors. Labels on dotted lines indicate cross-sectional differences. For example, “T < C, S” in (A) indicates that the temporal factor was associated with the worst baseline memory among AD dementia patients. Labels in the intervals indicate differences in longitudinal decline rates. For example, “ΔT, ΔC < ΔS” in (B) indicates that among Aβ- MCI participants, the temporal and cortical factors were associated with faster executive function decline than the subcortical factor. The schematics summarize the behavioral results of Figures 5, 6 and 7 (see **Supplemental Results** of **SI** for more discussion). Within each cognitive domain, the atrophy factors were associated with distinct trajectories across the stages. The trajectories of the cortical and subcortical factors transpose between the two cognitive domains. Divergence in memory trajectories existed even at the asymptomatic stage of the disease, i.e., among Aβ- CN participants

## Discussion

In this study, we identified distinct atrophy factors within AD dementia patients using Bayesian LDA modeling of MRI gray matter density maps. This approach estimated the factor composition of multiple atrophy factors for each participant rather than assuming membership to a single atrophy subtype (Fig. 1). Our analysis yielded a nested hierarchy of atrophy factors (Fig. 2), which corresponded to distinct trajectories of memory and executive function decline across the disease spectrum (Fig. 8). Overall, these results provide evidence that heterogeneity in patterns of atrophy exists in late-onset AD and that these atrophy patterns are associated with distinct cognitive trajectories.

### Atrophy Patterns in AD Dementia

Our model revealed a hierarchy of atrophy patterns within AD dementia patients (Fig. 2). As the number of estimated atrophy factors was increased from K to K+1, one atrophy pattern fractionated into two atrophy patterns, while the remaining patterns remained unchanged (Fig. S2). It is noteworthy that the atrophy patterns extracted using K=3 were similar to results from other groups investigating AD subtypes (Whitwell et al., 2012, Noh et al., 2014, Byun et al., 2015), although notable differences did emerge.

Specifically, our three-factor model revealed a “temporal” factor associated with atrophy in the temporal cortex, hippocampus and amygdala, a “cortical” pattern associated with atrophy in the frontal, parietal, lateral temporal and lateral occipital cerebral cortices, and a “subcortical” factor associated with atrophy in the cerebellum, striatum and thalamus (Fig. 2B). Our temporal factor was similar to the previously described “limbic-predominant” subtype, whereas the cortical factor was similar to the “hippocampal-sparing” subtype (Whitwell et al., 2012, Murray et al., 2011). More specifically, previous pathologically-defined subtypes were identified based on the ratio of NFT burden in hippocampal subregions versus association cortex, resulting in a limbic-predominant subtype and a hippocampal-sparing subtype. Follow-up VBM analyses (Whitwell et al., 2012) suggested gray matter loss in the temporoparietal cortex, frontal cortex, insula and precuneus in the hippocampal-sparing subtype, consistent with our cortical atrophy factor. On the other hand, Whitwell and colleagues identified predominant atrophy in the medial temporal lobe of the limbic-predominant subtype, consistent with our temporal atrophy factor.

A benefit of our approach is that the nested hierarchy of atrophy patterns was not mandated by our model but completely data-driven. Thus, although not mandated, our results revealed a nested hierarchy in contrast with previous approaches where hierarchy was imposed (e.g., Noh et al., 2014). Specifically, Noh and colleagues identified three subtypes: a “medial temporal” subtype, a “parietal frontal-dominant” subtype, and a “diffuse” subtype. Our temporal atrophy factor might correspond to their medial temporal subtype, whereas our cortical factor might correspond to their parietal frontal-dominant subtype, although direct comparison was difficult since their analyses were restricted to the cerebral cortex.

Our model suggests that atrophy patterns in AD patients follow a nested hierarchy structure. Given the nested hierarchy of cognitive functions revealed by a recent large-scale meta-analysis of ten thousand brain imaging experiments (Yeo et al., 2015) as well as brain network analyses (Zhou et al., 2006, Bassett et al., 2008, Meunier et al., 2009, Yeo et al., 2011), one might speculate that the nested hierarchy of atrophy factors arises from a natural hierarchy of brain functions and networks.

### Atrophy Factors Reflect Subtypes Rather Than Disease Stages

A potential pitfall of AD subtype analyses (Ritchie & Touchon, 1992) is that the observed heterogeneity might correspond to different disease stages (stage hypothesis), rather than heterogeneity in disease expression (subtype hypothesis). There are various reasons why the atrophy factors discussed in this manuscript likely correspond to subtypes rather than disease stages. First, there was not a single factor associated with the worst memory and executive function. Instead, decline trajectories of the temporal and cortical factors varied in their associations with the two cognitive domains (Fig. 8). Furthermore, analysis of follow-up MRI scans revealed that factor compositions were stable over time (Fig. 3), suggesting that individuals were not progressing from one factor to another, e.g., from temporal factor to cortical factor as predicted under the Braak staging scheme (Braak & Braak, 1991).

### Factor-Dependent Characteristics

There were significant differences across the atrophy factors in baseline age (p = 8e-7) and age at AD onset (p = 1e-5). Baseline age is dependent on study design, so drawing meaningful comparisons with the literature is difficult. Nevertheless, the cortical factor was associated with younger age at AD onset, consistent with previous studies describing subtypes with predominant cortical atrophy (Murray et al., 2011, Noh et al., 2014, Ossenkoppele et al., 2015). Importantly, years from AD onset to baseline did not differ across the three latent factors (p = 0.29; Table S2), providing further evidence that these factors were not simply disease stages. The subcortical factor was associated with a higher prevalence of the APOE ε2 allele (p = 0.03; not significant when corrected for multiple comparisons). The protective effects of the ε2 allele (Corder et al., 1994) might potentially contribute to the observation that the subcortical factor was associated with the mildest decline in both memory and executive function across all stages (Fig. 8).

Importantly, a lack of association between each factor and amyloid status suggests that atrophy factors do not merely reflect patterns associated with non-AD dementia patients that may have been “misdiagnosed” as AD dementia within the ADNI dataset (Lowe et al., 2013). However, even though repeating our factor estimation with Aβ+ AD dementia patients revealed consistent atrophy patterns with the model utilizing all AD patients, we are not able to determine whether atrophy patterns are a result of Aβ pathology or precede Aβ pathology. For instance, these atrophy patterns may emerge through processes not directly linked to Aβ pathology, but instead converge with AD pathology to influence disease progression. It is possible that factors such as co-morbid TDP-43 pathology, genetics, as well as development differences contribute to this heterogeneity. Along these lines, recent work suggests that different pathologies have distinct impacts on cognitive trajectories (Wilson et al., 2016). Interestingly, TDP-43 was shown to have a very early impact on cognitive trajectories compared to other pathologies such as hippocampal sclerosis and Lewy bodies. Given that TDP-43 is known to impact the medial temporal lobe (Josephs et al., 2014), it is possible that the temporal atrophy factor is influenced by the involvement of this pathology (since the temporal factor shows an early impact on memory among Aβ+ CN in our study).

A fundamental question that remains is why the expression of these atrophy patterns varies across individuals, especially since the spatial distribution of Aβ tends to be very diffuse throughout cortex. A similar dissociation is observed among AD patients with atypical clinical presentations, such that although the spatial pattern of Aβ is diffuse, the underlying pattern of NFTs and gray matter atrophy aligns with clinical symptoms (Ossenkoppele et al., 2016). Future work should investigate the time course of these atrophy patterns using longitudinal MRI as well as longitudinal assessment of Aβ and also investigate the prevalence of atrophy patterns among Aβ- participants to understand whether these patterns are specific for AD or merely converge with AD processes to influence disease progression.

### Distinct Memory and Executive Function Decline Trajectories

The behavioral results (Figs. 5, 6 and 7) are summarized in Fig. 8. Overall, we found that the associations between atrophy factors and cognition varied by domain, as well as time course in the disease. Specifically, the temporal factor showed the greatest association with memory, a relationship that emerged early among Aβ+ CN participants and remained consistent in later disease stages. Conversely, the cortical factor was associated with both memory and executive function, but exerted greater impact later in the disease among Aβ+ MCI participants and AD patients.

Overall, the trajectories (Fig. 8) revealed several salient points. First, memory decline in the context of late-onset AD occurred earlier than decline in executive function, which is in line with previous studies (Grober et al., 2008). Second, divergence of memory trajectories among atrophy factors Appeared as early as the asymptomatic (CN) stage of the disease, whereas divergence of executive function trajectories was not detectable until the MCI stage (Fig. 8). Specifically, the temporal and subcortical factors showed faster memory decline than the cortical factor among Aβ+ CN participants, and by MCI, the temporal factor was already associated with worse memory at baseline than the subcortical factor. In contrast, there was no difference in executive function decline rates among Aβ+ CN participants or cross-sectional differences among Aβ+ MCI participants. Interestingly, AD dementia patients expressing the cortical factor exhibited the fastest decline rates in both executive function and memory. Finally, the subcortical factor (blue curves in Fig. 8) was the mildest factor in terms of both memory and executive function deterioration. In both Aβ+ MCI and AD dementia participants, the subcortical factor was associated with the best memory and executive function scores, as well as the slowest decline rates.

### Correspondence and Extensions of AD Heterogeneity Literature

Our results were consistent with the preponderance of literature on heterogeneity among AD dementia patients. For example, our atrophy factors show overlap with the pathologically-defined hippocampal-sparing and limbic-predominant subtypes (Whitwell et al., 2012), as well as the subtypes described by Noh et al. (2014). Our analyses suggested that the cortical factor was associated with faster decline in both memory and executive function than the temporal factor at the dementia stage, which is consistent with the hippocampal-sparing subtype exhibiting faster MMSE decline than the limbic-predominant subtype among AD dementia patients (Murray et al., 2011). Similarly, our finding that the cortical factor was associated with the most rapid memory and executive function decline among AD dementia patients was also consistent with Byun et al. (2015). Among AD dementia patients, the cortical factor was associated with the worst baseline executive function, while the temporal factor was associated with the worst baseline memory. This is consistent with previous work showing that thinning of frontoparietal cortical regions was associated with nonamnestic presentations and dysexecutive phenotypes (Dickerson & Wolk, 2011) and that the “cortical atrophy-only” subtype had worse baseline executive function than the “hippocampal atrophy-only” subtype (Byun et al., 2015). Thus, our data-driven approach provides further evidence that distinct atrophy patterns among AD patients impact different cognitive domains.

In addition to characterizing heterogeneity among AD dementia patients, we extended our approach to participants that were presumably in very early stages of AD development (i.e., Aβ+ but without the clinical symptoms of dementia; Sperling et al., 2011, Albert et al., 2011). By examining earlier stages, we found that the temporal factor showed the greatest association with memory decline among Aβ+ CN participants, but that the cortical factor was a stronger predictor of memory decline among AD dementia patients (Fig. S7A2). Likewise, although the cortical factor was not associated with either cognitive domain among Aβ+ CN participants, this factor was associated with executive function decline in Aβ+ MCI participants and AD patients (Figs. 7B and 8). The impact of these atrophy factors at different points along the clinical spectrum has important implications for measuring decline and understanding the progression of AD. Furthermore, consideration of this heterogeneity may improve the ability to identify individuals most at-risk for cognitive decline as compared to approaches that measure atrophy using the same regional metric across all participants.

### Mixed Membership Modeling and Precision Medicine

One key advantage of our modeling strategy is that individuals can express multiple latent atrophy factors (i.e., mixed membership) rather than being assigned to a single subtype. Therefore, patients classified by Murray et al. (2011) as hippocampal-sparing (or limbic-predominant) might correspond to the few patients in our study that predominantly expressed the cortical (or temporal) atrophy factor. Murray et al. (2011) defined a third group of patients that were considered “typical” by virtue of being neither hippocampal-sparing nor limbic-predominant. These typical patients might correspond to the majority of AD dementia patients in our study that expressed multiple latent factors to similar degrees.

The use of mixed membership modeling has implications for estimation of factor-dependent atrophy maps and cognitive decline. For example, consider a hypothetical patient who expressed 50% subcortical, 40% temporal and 10% cortical factors. In our analyses, 50%, 40% and 10% of the patient’s atrophy map would contribute to the estimation of the probabilistic atrophy maps of the subcortical, temporal and cortical factors, respectively. This extends previous approaches (Whitwell et al., 2012, Noh et al., 2014, Byun et al., 2015) that classified each patient into one single subtype and then performed group comparisons to obtain differential atrophy patterns, despite the fact that each patient might express multiple latent atrophy factors. Thus, more information about each participant is retained by treating factor compositions continuously rather than assigning participants to a single group.

Similarly, 50%, 40% and 10% of the hypothetical patient’s cognitive decline rate would contribute to our estimation of the memory decline rates associated with the subcortical, temporal and cortical factors, respectively. Indeed, when such a patient was simply assigned to a single factor based on the highest probability (i.e., assigned to a pure subtype), the estimated differences in cognitive decline rates across subtypes were found to be substantially weaker. The reason should be clear when considering the hypothetical patient. Since the patient expressed 50% subcortical, 40% temporal and 10% cortical factors, one would expect the memory decline rate to be faster than a pure subcortical subtype (and slower than a pure temporal factor). By assigning this patient to be a pure subcortical subtype, one would overestimate the decline rate of the subcortical subtype.

Although we observe some participants with extreme probabilities of a single atrophy factor, these participants are infrequent. Instead, the majority of the participants expressed intermediate probabilities across multiple latent atrophy factors. We can potentially utilize the factor decomposition to predict the memory and executive function decline trajectories of individual participants. For example, we might predict the hypothetical patient who expressed 50% subcortical, 40% temporal and 10% cortical factors to have decline trajectories corresponding to 50% times the blue curve plus 40% times the green curve plus 10% times the red curve from Fig. 8. Therefore, the factor composition can be thought of as an individualized subtype diagnosis of the participant, representing a small but crucial step towards precision medicine.

### Limitations

Our study has multiple limitations. First, direct comparisons with other subtype studies were difficult because of methodological differences, including the utilization of mixed membership modeling and participant selection. Another limitation is the arbitrary choice of the number of latent atrophy factors to estimate using LDA. Given consistency with previous studies and a limited sample size, we focused on K = 2 to 4 factors, but atrophy factors beyond K=4 may be biologically relevant.

### Conclusion

By utilizing a novel Bayesian modeling framework, our study revealed three latent AD atrophy factors with distinct memory and executive function trajectories. Across the clinical spectrum, the cortical atrophy factor was associated with the worst executive function performance, while the temporal atrophy factor was associated with worst memory performance. The subcortical atrophy factor has not been discussed in the literature and was associated with the slowest memory and executive function decline. Our approach allowed each individual to express multiple atrophy factors to various degrees rather than assigning the individual to a single subtype. Therefore, each participant exhibited his or her own unique factor composition, which can potentially be exploited to predict individual-specific cognitive decline trajectories, with potential implications for prevention and monitoring disease progression. Finally, our methodological framework is general and can be utilized to discover subtypes in other brain disorders. Code from this manuscript is publicly available at link_to_be_added.

## Materials and Methods

**Overview.** Voxelwise atrophy of 188 AD dementia patients was derived from their structural MRI data (Ashburner & Friston, 2000, Douaud et al., 2007). Subsequent analyses proceeded in three steps. In **Step I**, a Bayesian model (Fig. 1; Blei et al., 2003) was applied to estimate the probabilistic atrophy maps of latent factors Pr(Voxel | Factor) and the factor composition of each patient Pr(Factor | Patient). The probabilistic atrophy maps were then used to infer the factor compositions of 43 Aβ+ CN participants and 147 Aβ+ MCI participants. In **Step II**, stability of the factor decomposition over a period of two years was analyzed. In addition, characteristics (demographics, age at AD onset, years from AD onset to baseline, amyloid burden and APOE genotype) of all participants were compared across the factors. Finally, in **Step III**, we analyzed the atrophy factors' relationships with cross-sectional baseline and longitudinal decline of memory and executive function. Each step is described in detail below.

**Data.** Data used in this study were obtained from the Alzheimer’s Disease Neuroimaging Initiative (ADNI) database (http://adni.loni.usc.edu), launched in 2003 as a public-private partnership and led by Principal Investigator Michael W. Weiner, MD. The primary goal of ADNI has been to test whether serial MRI, positron emission tomography, other biological markers, and clinical and neuropsychological assessment can be combined to measure the progression of MCI and early AD (for up-to-date information, see http://www.adni-info.org/). Institutional review boards approved study procedures across participating institutions (see **SI** for the complete list of the institutions). Written informed consent was obtained from all participants.

This study considered the structural MRI (T1-weighted, 1.5 Tesla) of 810 participants enrolled in ADNI 1, comprising 188 AD dementia (at baseline, same hereinafter) patients, 394 MCI participants and 228 CN participants. Of the 188 AD dementia patients, 100 had their CSF amyloid data available, and 91 of the 100 were Aβ+. AD onset was on average 3.6 years (std = 2.5, min = 0, max = 13) before baseline. Of the 394 MCI participants, 197 had their CSF amyloid data available, and 147 of the 197 were Aβ+. Of the 228 CN participants, 114 had their CSF amyloid data available, and 43 of the 114 were Aβ+. The Aβ+ CN elderly participants and the Aβ+ MCI participants are referred to as the Aβ+ nondemented group (N = 190) in this study.

According to the ADNI protocol, AD dementia patients had their cognition examined at baseline, in months 6, 12 and 24. In addition, normal participants were examined in month 36 and annually afterwards. MCI participants underwent another extra exam in month 18. Although this study only considered participants enrolled in ADNI 1, to increase statistical power, their neuropsychological scores (ADNI-Mem, ADNI-EF and MMSE) from ADNI GO and ADNI 2 were also included in the longitudinal analyses of cognitive decline.

**Voxel-Based Morphometry.** Structural MRI data of all 810 participants were analyzed with FSL-VBM (http://fsl.fmrib.ox.ac.uk/fsl/fslwiki/FSLVBM; Douaud et al., 2007), a VBM protocol (Good et al., 2001) carried out with FSL tools (Smith et al., 2004). First, structural images were brain-extracted and gray matter (GM) segmented before being registered to the MNI152 standard space using affine registration. Second, the affine-registered images were flipped about the x-axis and averaged to create a left-right symmetric, study-specific affine GM template. Third, the GM images were nonlinearly registered to the affine GM template and were again flipped and averaged to create a final left-right symmetric, study-specific nonlinear GM template in MNI152 space. Fourth, all native GM images were nonlinearly registered to this final template and modulated to account for local expansion (or contraction) due to the nonlinear component of the spatial transformation. The resulting GM density images were smoothed with a Gaussian kernel of 10mm full width at half maximum (FWHM), consistent with standard VBM practices (e.g., Dole et al., 2013, Pardoe et al., 2008). Finally, we applied log10 to the smoothed GM density images and regressed out possible effects of age, sex and intracranial volume (ICV) with a general linear model (GLM) estimated from just the 228 CN participants.

**Quality Control for Voxel-Based Morphometry.** The outputs of each VBM step were visually checked by authors XZ and NS. Details are found in **Supplemental Methods** of **SI**.

**Bayesian Model.** We sought a mathematical model that captured the premise that each AD patient expresses one or more latent atrophy factors, each of which is associated with distinct, but possibly overlapping atrophy patterns (Fig. 1). Among many possible models, the latent Dirichlet allocation (LDA) model (Blei et al., 2003) is probably the simplest and was applied to the ADNI data.

The LDA model was originally developed to automatically discover latent topics in a collection of text documents. The model assumes that each document is an unordered collection of words associated with a subset of K latent topics. Each topic is represented by a probability distribution over a dictionary of words. Given a collection of documents, there exist algorithms (Blei et al., 2003) to estimate the probability of a dictionary word given a topic Pr(Word | Topic) and the probability that a topic is associated with a particular document Pr(Topic | Document). The LDA model is useful because it allows a document to be associated with multiple topics (which can be shared across documents) and each topic to be associated with multiple words (which can be shared across topics).

To map the LDA model to the ADNI data, one can think of AD patients as text documents, atrophy factors as topics, and MNI152 voxels as dictionary words. Correspondingly, each patient expresses one or more latent atrophy factors to different extents (Pr(Factor | Patient)), and each factor is associated with atrophy at multiple voxels to different extents (Pr(Voxel | Factor)).

LDA assumes that a document is summarized by the number of times a dictionary word appears in the document. Since dictionary words correspond to MNI voxels, the continuous log-transformed GM density images (previous section) were discretized so that greater atrophy corresponded to larger word counts. More specifically, for each voxel of the log-transformed GM density images, z-transformation (with respect to the 228 CN participants) was performed for each of the 810 participants. Therefore, a z-score of less than 0 at a given voxel of a particular individual would imply above-average atrophy at the voxel relative to the CN participants. Z-scores above zero were set to 0, equivalent to regarding the voxels as atrophy-free. Finally, the z-scores were multiplied by-10 and rounded to the nearest integer, so that larger positive values (greater word count) indicated more severe atrophy.

The LDA model assumes that the ordering of words within a document is exchangeable. In the context of our application, the corresponding assumption is that the ordering of atrophied voxels is exchangeable. Although word order in real documents is important, the ordering of atrophied regions (e.g., prefrontal vs. parietal) reported in an experiment is arbitrary and thus consistent with the assumption. Consequently, the LDA model appears particularly well suited for applications in the present context.

Given the discretized voxelwise atrophy of the 188 AD dementia patients and the number of latent atrophy factors K, the variational expectation-maximization (VEM) algorithm (http://www.cs.princeton.edu/~blei/lda-c/; Blei et al., 2003) was applied to estimate Pr(Factor | Patient) and Pr(Voxel | Factor). For each K, the algorithm was rerun with forty different random initializations, and the solution with the highest likelihood (bound) was selected. The random initializations led to highly similar solutions, suggesting that forty random initializations were sufficient for robust factor estimations.

The probabilistic atrophy maps Pr(Voxel | Factor) estimated from the AD dementia patients were used to infer factor compositions Pr(Factor | Participant) of the 190 Aβ+ nondemented participants using the standard VEM algorithm (Blei et al., 2003).

**Interpreting Pr(Voxel | Factor) and Pr(Factor | Patient).** For a given latent factor, Pr(Voxel | Factor) is a probability distribution over all the GM voxels, which can be visualized as a probabilistic atrophy map overlaid on the FSL MNI152 template (each row of Fig. 2).

Pr(Factor | Patient) is a probability distribution over latent atrophy factors, representing the factor composition of the patient, and can be visualized as a dot inside a “factor triangle” (for K=3 factors) whose barycentric coordinates equal Pr(Factor | Patient) as shown in Figs. 4 and S5A. For example, Pr(Factor | Patient) = [0.7, 0.2, 0.1] implies that the patient expresses a pattern of brain atrophy due to 70% temporal, 20% subcortical and 10% cortical factors and that the dot representing this patient falls closer to the “temporal corner” of the factor triangle. This contrasts with work in the literature that assigns each individual to a single subtype (e.g., Murray et al., 2011, Noh et al., 2014, Byun et al., 2015).

**Quantifying the Nested Hierarchy of Atrophy Factors.** An important model parameter is the number of latent factors K. Therefore, we determined how factor estimation changed from K = 2 to 10 factors. See **Supplemental Methods** of **SI** for the detailed description.

**Top Anatomical Structures Associated with Each Factor.** This manuscript focuses on three atrophy factors. To automatically identify the gray matter anatomical structures most associated with each atrophy factor, the MNI152 template was first processed using FreeSurfer 4.5.0 (Fischl, 2012). The FreeSurfer software automatically segmented the MNI152 template into multiple cortical (Fischl et al., 2004, Desikan et al., 2006) and subcortical (Fischl et al., 2002, Fischl et al., 2004) structures, such as the inferior parietal cortex and hippocampus. For each anatomical structure, we averaged Pr(Voxel | Factor) over all its voxels. The structure was assigned to the factor with the largest average probability. For each factor, we tabulated the assigned brain structures and ranked them in the descending order of average probability. The results are in Table S1.

**Cross-Pipeline Validation of Atrophy Patterns.** To ensure the atrophy factors were robust to choice of VBM software (FSL), we performed posthoc analyses using FreeSurfer. Details are found in **Supplemental Results** and **Supplemental Methods** of **SI**.

**Atrophy Factor Stability.** To examine the atrophy factor stability during disease progression, we considered all 810 participants who had their two-year follow-up scans available (N = 560). First, their baseline factor compositions Pr(Factor | Participant) were extracted using their baseline MRI data. Next, VBM was performed on the follow-up structural MRI data using the VBM template previously created with all 810 participants. Subsequent processing (e.g., z-normalization) adopted parameters used in processing the 810 baseline scans. Factor compositions were then inferred with the processed VBM results (same procedure as inferring factor compositions of Aβ+ CN and MCI participants). The factor stability was visualized with a scatter plot for each factor (Figs. 3 and S4). Each participant is represented by a dot whose x-coordinate is the factor composition at baseline and y-coordinate is the factor composition after two years. Therefore, if the factor estimation is stable over disease progression, one would expect a close-to-one correlation coefficient and a y = x linear fit.

**Comparing Patient Characteristics by Atrophy Factor.** We explored how patient characteristics (baseline age, age at AD onset, years from onset to baseline, education, sex, amyloid and APOE genotype) varied across the three latent factors (Table S2) using GLM (and logistic regression for binary variables).

GLM was applied to baseline age, age at AD onset, years from onset to baseline, education, amyloid and APOE: the characteristic of interest served as response y, and the subcortical factor probability s and cortical factor probability c were included as explanatory variables. Hence, the GLM was y=β_0_+β_s_·s+β_c_·c+ε, where β’s are the regression coefficients, and ε is the residual. The temporal factor probability t was implicitly modeled because t + s + c = 1. Intuitively, β_0_ reflected the response of the temporal factor, β_s_ reflected the response difference between the subcortical and temporal factors, and β_c_ reflected the difference between the cortical and temporal factors.

Statistical tests of whether the characteristic y varied across factors involved null hypotheses of the form Hβ = 0, where β=[β_0_, β_s_, β_c_]^T^, and H is the linear contrast (Koch, 1999). We first performed a statistical test of overall differences across all factors with H=[0, 1, 0; 0, 0, 1]. We then tested for differences between the factors. For example, H=[0, 1, −1] tested possible differences between the subcortical and cortical factors. H=[0, 1, 0] compared the subcortical and temporal factors. Similarly, H=[0, 0, 1] compared the cortical and temporal factors.

Since sex is a binary variable, logistic regression was applied. In this case, response y was sex (0 for male, 1 for female), and explanatory variables consisted of the subcortical factor probability s and cortical factor probability c. Therefore, the regression model was log(μ/(1-μ))=β_0_ + β_s_·.s + β_c_·c + ε, where μ is the probability of female, β’s are the regression coefficients, and ε is the residual. Intuitively, the linear combination β_0_ + β_s_·s + β_c_·c predicts the probability of female (y = 1). exp(β_0_) reflects the odds ratio for the temporal factor; exp(β_s_) reflects the ratio of odds ratio between the subcortical and temporal factors; exp(β_c_) reflects the ratio of odds ratio between the cortical and temporal factors.

Likelihood ratio test was utilized to determine whether sex varied across the latent atrophy factors. In short, the test involved comparing the likelihood of an appropriately restricted model to the original model (Koch, 1999). We first performed a statistical test of overall differences across factors. In this case, the restricted model log(μ/(1-μ))=β_0_ + ε was fitted to the data, and the resulting likelihood was compared with the likelihood of the original model y =p_0_ + β_s_·s + β_c_·c + ε. We then tested for possible differences between atrophy factors. For example, to compare the subcortical and cortical factors, the restricted model was log(μ/(1-μ)) = β_0_ + β_s_·(s+c) + ε because β_s_ = β_c_ under the null hypothesis. To compare the subcortical and temporal factors, the restricted model became log(μ/(1-μ)) = β_0_ + β_c_·c + ε because β_s_ = 0 under the null hypothesis. To compare the cortical and temporal factors, the restricted model was log(μ/(1-μ)) = β_0_ + β_s_·s + ε because β_c_ = 0 under the null hypothesis.

**General Linear Modeling of Cross-sectional Cognition Among Aβ+ CN, Aβ+ MCI and AD Dementia Participants.** A single GLM was utilized to examine cross-sectional differences in memory (ADNI-Mem; Crane et al., 2012) across the atrophy factors in the 43 Aβ+ CN, 147 Aβ+ MCI, and 188 AD dementia participants. The same model was estimated for K = 2, 3 and 4 factors, as well as for executive function (ADNI-EF; Gibbons et al., 2012) and MMSE.

For ease of explanation, we will focus on explaining the GLM for the case of three atrophy factors and ADNI-Mem. Response y of the GLM consisted of the 378 (= 43 CN + 147 MCI + 188 AD) participants’ baseline ADNI-Mem. Explanatory variables consisted of binary MCI group indicator m, binary AD dementia group indicator d, subcortical factor probability s, cortical factor probability c, and interactions between group indicators and factor probabilities (i.e., m·s, m·c, d·s and d·c), while nuisance variables consisted of baseline age x_1_, sex x_2_, education x_3_ and total atrophy x_4_ (defined as ICV divided by total GM volume as estimated by FSL).

Therefore, the GLM was y = β_0_ + β_m_·m + β_d_·d + β _s_·s + β_c_·c + β_ms_·m·s + β_mc_·mc + β_dc_·d·s + β_dc_dc + β_1_·x_1_ + β_2_x_2_ + β_3_x_3_ + β_4_x_4_ + ε, where β’s are the regression coefficients, and ε is the residual. Temporal factor probability t was implicitly modeled because t + s + c = 1. This was also the case for CN group indicator n because only one of n, m and d is 1 with the other two being 0. Intuitively, β_0_ reflected the temporal factor’s contribution to ADNI-Mem at the CN baseline (because m = d = s = c = 0), β_0_ + β_m_ reflected the temporal factor’s contribution to ADNI-Mem at the MCI baseline (because m = 1, and d = s = c = 0), and β_0_ + β_m_ + β_s_ + β_ms_ reflected the subcortical factor’s contribution to ADNI-Mem at the MCI baseline (because m = 1, s = 1, and d = c = 0). With this model setup, variations in age, sex, education and total atrophy were controlled for across participants.

Statistical tests involved null hypotheses of the form Hβ = 0, where β = [β_0_, β_m_, β_d_, β_s_, β_c_, β_ms_, β_mc_, β_ds_, β_dc_, β_1_, β_2_, β_3_, β_4_]^T^, and H is the linear contrast (Koch, 1999). First, we tested whether ADNI-Mem deteriorated across disease stages (i.e., from CN to MCI to AD) for each factor. Specifically, for each factor, we tested possible differences in ADNI-Mem between the CN and MCI baselines, MCI and AD baselines, and CN and AD baselines. For example, to test whether ADNI-Mem deteriorated significantly from the CN to MCI baseline for the temporal factor, H was specified such that Hβ = β_m_ = 0. As another example, Hβ = β_d_ + β_dc_−β_m_−β_mc_ = 0 tested whether ADNI-Mem degraded greatly from the MCI to AD baseline for the cortical factor. The test results for both memory and executive function are tabulated in Figs. 5A1 and 5B1.

To foreshadow the results, the hypothesis tests in the previous paragraph hinted at differences in cross-sectional ADNI-Mem across the factors. Therefore, statistical tests of whether cross-sectional ADNI-Mem y varied across factors at each disease stage were performed. For each stage baseline, we first performed a statistical test of overall differences across all factors and then tested for pairwise differences. Take the AD baseline as an example. To test whether baseline memory differed across all factors among AD dementia patients, H was specified such that Hβ = 0 translated to β_s_ + β_ds_ = β_c_ + β_dc_ = 0. For pairwise comparisons, β_s_ + β_ds_ = 0 tested possible differences between the temporal and subcortical factors at the AD baseline, β_c_ + β_dc_ = 0 compared the temporal and cortical factors at the AD baseline, and β_s_ + β_ds_ = β_c_ + β_dc_ tested possible differences between the subcortical and cortical factors at the AD baseline.

The results of the above statistical tests are shown in Figs. 5A1, 5B1, 6, S6A1, S6B1, S7A1, S7B1, S7C1, S8A1 and S8B1, where (except in Figs. 5A1 and 5B1) the blue dot corresponds to the estimated difference in baseline scores between two “pure factors” after controlling for age, sex, education and total atrophy. For example, when comparing subcortical and cortical factors at the MCI baseline, the estimated difference in baseline cognition is given by β_s_ + β_ms_−β_c_−β_mc_. The red bar corresponds to the standard error of this estimation given by std(β_s_ + β_ms_−β_c_−β_mc_).

**Linear Mixed-Effects Modeling of Longitudinal Cognitive Decline Among Aβ+ CN, Aβ+ MCI and AD Dementia Participants.** To analyze variations in cognitive decline rates across atrophy factors, we utilized the linear mixed-effects (LME) model, whose setup was similar to the GLM setup (previous section). Details are found in **Supplemental Methods** of **SI**. Results of the LME statistical tests are illustrated in Figs. 5A2, 5B2, 7, S6A2, S6B2, S7A2, S7B2, S7C2, S8A2 and S8B2.

**False Discovery Rate Correction for Behavioral Tests.** Because of the many statistical tests performed in the behavioral analyses, multiple testing was corrected using false discovery rate (FDR; Benjamini & Hochberg, 1995) at q = 0.05 for all behavioral comparisons. In detail, included tests are diagnostic group comparisons in memory and executive function regardless of factors as well as all comparisons of baseline and longitudinal decline rates of memory, executive function and MMSE at all disease stages for K = 2, 3 and 4 factors. In total, we corrected for 240 statistical tests. P values that remained significant after FDR control were highlighted in blue in Figs. 5, 6, 7, S6, S7 and S8.

## Acknowledgements

This work was supported by NUS Tier 1, Singapore MOE Tier 2 (MOE2014-T2-2-016), NUS Strategic Research (DPRT/944/09/14), NUS SOM Aspiration Fund (R185000271720), Singapore NMRC (CBRG14nov007, NMRC/CG/013/2013) and NUS YIA. The research also utilized resources provided by the Center for Functional Neuroimaging Technologies, P41EB015896 and instruments supported by 1S10RR023401, 1S10RR019307, and 1S10RR023043 from the Athinoula A. Martinos Center for Biomedical Imaging at the Massachusetts General Hospital. Data collection and sharing for this project was funded by the Alzheimer's Disease Neuroimaging Initiative (ADNI) (National Institutes of Health Grant U01 AG024904) and DOD ADNI (Department of Defense award number W81XWH-12-2-0012). ADNI is funded by the National Institute on Aging, the National Institute of Biomedical Imaging and Bioengineering, and through generous contributions from the following: AbbVie, Alzheimer's Association; Alzheimer's Drug Discovery Foundation; Araclon Biotech; BioClinica, Inc.; Biogen; Bristol-Myers Squibb Company; CereSpir, Inc.; Eisai Inc.; Elan Pharmaceuticals, Inc.; Eli Lilly and Company; EuroImmun; F. Hoffmann-La Roche Ltd and its affiliated company Genentech, Inc.; Fujirebio; GE Healthcare; IXICO Ltd.; Janssen Alzheimer Immunotherapy Research & Development, LLC.; Johnson & Johnson Pharmaceutical Research & Development LLC.; Lumosity; Lundbeck; Merck & Co., Inc.; Meso Scale Diagnostics, LLC.; NeuroRx Research; Neurotrack Technologies; Novartis Pharmaceuticals Corporation; Pfizer Inc.; Piramal Imaging; Servier; Takeda Pharmaceutical Company; and Transition Therapeutics. The Canadian Institutes of Health Research is providing funds to support ADNI clinical sites in Canada. Private sector contributions are facilitated by the Foundation for the National Institutes of Health (www.fnih.org). The grantee organization is the Northern California Institute for Research and Education, and the study is coordinated by the Alzheimer's Disease Cooperative Study at the University of California, San Diego. ADNI data are disseminated by the Laboratory for Neuro Imaging at the University of Southern California.

## Supporting Information

This supplemental material is divided into **Supplemental Results**, **Supplemental Methods**, **Supplemental Figures and Tables**, and **Complete List of ADNI Investigators and Participating Institutions.**

### Supplemental Results

#### Similar Atrophy Factors Were Obtained from Aβ+ MCI Participants

We confirmed that atrophy patterns estimated with our LDA approach would be similar during the nondemented stage compared to the resulting factors from the AD dementia group. Given the small number of the Aβ+ CN participants, we estimated atrophy factors with the 147 Aβ+ MCI participants (Fig. S3C) and confirmed that the obtained atrophy factors were highly similar, with an average correlation across all pairwise comparisons of r = 0.77. Therefore, the atrophy factors from the AD dementia patients were utilized for subsequent analyses.

#### Atrophy Factors Were Robust to Choice of Software

Table S1 lists the anatomical structures associated with each factor based on overlap between the atrophy maps and anatomical structures in MNI152 space as defined by FreeSurfer [1] (see **Supplemental Methods**). The volumes of individual anatomical structures in all AD dementia patients were computed using FreeSurfer. Regression analyses confirmed that volumes of anatomical structures associated with an atrophy factor were lower (after controlling for intracranial volume) in participants with higher loading on the factor (see **Supplemental Methods**). For example, the temporal factor was associated with the most severe atrophy in the structures listed by Table S1A compared with the subcortical factor (p = 2e-15) and cortical factor (p = 4e-15), whereas there were no differences between the subcortical and cortical factors (p = 0.84). Results for the subcortical and cortical factors are in the captions of Tables S1B and S1C. The agreement between FSL-VBM [2] and this posthoc analysis with FreeSurfer suggested that the factors were unlikely the results of segmentation or registration artifacts.

#### Baseline and Longitudinal Decline of Memory and Executive Function Were Consistent Across Factor Hierarchy

The behavioral (memory and executive function) analyses were repeated for two and four atrophy factors (Figs. S6 and S8). The results were consistent with the hierarchy of atrophy factors.

For example, the temporal and subcortical factors in the three-factor model were merged as a single temporal+subcortical factor in the two-factor model. Since the cortical factor was associated with the fastest longitudinal memory decline among the three factors in the AD dementia cohort (Fig. 7A), we expected the cortical factor to be associated with faster memory decline than the temporal+subcortical factor in the two-factor model, which was indeed the case (p = 2e-6; Fig. S6A2).

On the other hand, the three-factor analysis of AD dementia patients suggested that the temporal factor was associated with worse memory than the cortical factor, while the cortical factor was associated with slightly worse memory than the subcortical factor (Fig. 6A). Therefore, we expected difference in baseline memory between the temporal+subcortical and cortical factors (in the two-factor model) to be diluted by the fusion of the temporal and subcortical factors, which was indeed the case (p = 0.17; Fig. S6A1). Therefore, additional insights into factor differences could be obtained by going from two factors to three factors.

As the number of factors was increased from three to four, the cortical factor split into frontal and posterior cortical factors. There was again consistency when comparing the four-factor results with the three-factor results. The two factors were mostly associated with similar behavioral trajectories, except that among Aβ+ MCI participants, the posterior cortical factor was associated with faster memory (p = 8e-3) and executive function (p = 9e-8) decline rates than the frontal cortical factor (Fig. S8).

As the number of factors increased, the effective (average) number of participants per factor decreased (e.g., the effective number of Aβ+ CN participants “assigned to the temporal factor” is only 5.7 for the four-factor model), thus reducing our confidence in larger number of factors despite the successful behavioral dissociation. Therefore, this work focused on interpreting the results of the three-factor model. As the ADNI database continues to grow, future work might re-visit the question of larger number of atrophy factors.

#### Schematics of Memory and Executive Function Trajectories Based on Statistical Test Results

The behavioral results (Figs. 5, 6 and 7) are summarized by the schematics of trajectories in Fig. 8, which were drawn based on how memory (or executive function) of each factor declined across disease stages and how the factors compared with each other in terms of memory (or executive function) decline at each stage.

All salient features of the trajectories reflect the results of statistical tests (Figs. 5, 6 and 7). For example, the executive function trajectories of all three atrophy factors were almost flat and did not diverge at the CN stage (Fig. 8B). This was based on the fact that there was no change in ADNI-EF [3] performance between Aβ+ CN and MCI participants for all three factors (Fig. 5B1), as well as no difference in ADNI-EF decline rates between factors among Aβ+ CN participants (Fig. 7B). From the MCI stage onwards, the trajectory of the cortical factor (red curve) became increasingly steep, reflecting the test results that executive function decline of the cortical factor accelerated from CN to MCI to AD (Fig. 5B2). This was also consistent with the ADNI-EF decrease between MCI and AD (Fig. 5B1). In contrast, trajectories of the temporal and subcortical factors (blue and green curves) remained almost flat from MCI to AD because there was no difference in ADNI-EF performance between MCI and AD for the two factors (Fig. S5B1). In addition, cross-sectional and longitudinal differences between the factors (Figs. 6B and 7B) were also respected in Fig. 8B, e.g., the cortical factor was associated with the worst baseline ADNI-EF and the most rapid decline among AD dementia patients.

One salient feature of the memory trajectories was the crossing of the subcortical and cortical factors (blue and red curves), supported by the following behavioral tests. Among Aβ+ CN participants, both the temporal and subcortical factors exhibited significant memory decline rates, but not the cortical factor (Fig. 5A2). The temporal and subcortical factors showed faster memory decline than the cortical factor (Fig. 7A). These results implied that the cortical (red) curve should be above the subcortical (blue) and temporal (green) curves immediately after CN (Fig. 8A). Among Aβ+ MCI participants, the temporal factor was associated with worse memory than the subcortical factor, but not the cortical factor (Fig. S7A1). This implies that the cortical (red) curve should be lower than the subcortical (blue) curve, closer to the temporal (green) curve. This is also consistent with the statistical test showing a significant decrease in memory performance between MCI and CN for the cortical and temporal factors, but not for the subcortical factor (Fig. 5A1). Together, the results imply that the cortical (red) curve, originally higher than the subcortical (blue) curve at the CN stage, later crossed the subcortical (blue) curve before the MCI stage.

### Supplemental Methods

**Quality Control for Voxel-Based Morphometry.** The outputs of each VBM step were visually checked by authors XZ and NS. In practice, all the VBM steps (except for brain extraction) did not require any manual interventions. The brain extraction (FSL BET [4]) sometimes resulted in inaccurate brain extraction, e.g., part of the neck was sometimes included as part of the brain. For these problematic cases, the parameters were manually tuned until the results were satisfactory. The 810 baseline scans and 560 follow-up scans (see the second paragraph of **II. Examining Factor Robustness and Characteristics of Factor Compositions**) were processed jointly to avoid bias introduced by processing the baseline and follow-up scans separately as two independent sets. Specifically, the 810 baseline scans and 560 follow-up scans were mixed together and randomly divided into two sets, such that each set contained both baseline and follow-up scans. XZ and NS each processed one set. To ensure common quality control standards, XZ and NS independently processed a small number of the participants, compared their conclusions, and eventually reached consensus.

**Quantifying the Nested Hierarchy of Atrophy Factors.** An important model parameter is the number of latent factors K. Therefore, we determined how factor estimation changed from K = 2 to 10 factors. An exhaustive search was performed to quantify the possibility that two atrophy patterns in the (K+1)-factor model were subdivisions of a pattern in the K-factor model (while the remaining K-1 atrophy patterns remained similar across both models). This quantification is based on the following idea: suppose an atrophy pattern in the K-factor model divides into the i-th and j-th patterns in the (K+1)- factor model, then the average of the i-th and j-th patterns should be similar to the original pattern. To quantify the presence of this phenomenon, the Pr(Voxel | Factor) of the i-th and j-th latent factors were averaged into a single Pr(Voxel | Factor). The resulting K factors of the (K+1)-factor model were matched to the K-factor model by reordering the factors (using the Hungarian matching algorithm) to maximize the correlation of Pr(Voxel | Factor) between corresponding pairs of factors. After obtaining the optimal correspondence, the pairwise correlations were averaged across all pairs of factors, resulting in an average correlation value indicating the quality of the split (with higher correlation values indicating a better split). By performing an exhaustive search over all pairs of i and j, we found the atrophy factor of the K-factor model whose split best approximated the (K+1)-factor model (Fig. S2A). This procedure was independently repeated using Pr(Factor | Patient) (Fig. S2B).

**Cross-Pipeline Validation of Atrophy Patterns.** To ensure the atrophy factors were robust to choice of VBM software (FSL [2]), we performed posthoc analyses using FreeSurfer. Recall from **Top Anatomical Structures Associated with Each Factor**, that we have assigned each MNI GM anatomical structure to each of the three atrophy factors (Table S1). The structural MRI data of the 378 (= 43 CN + 147 MCI + 188 AD) participants were preprocessed using FreeSurfer so as to obtain volume estimates of all the anatomical structures for each participant. We then verified using GLM that each factor had a smaller total volume of its assigned GM anatomical structures than the other two factors (while controlling for ICV).

For example, Table S1A shows the top GM anatomical structures associated with the temporal factor. A GLM was set up where the response variable y was the total volume of the anatomical structures listed in Table S1A, while the explanatory variables included the subcortical factor probability s, cortical factor probability c, and ICV i. Hence, the GLM was y = β_0_ + β_s_·s + β_c_·c + β_i_·i + ε, where β’s are the regression coefficients, and ε is the residual. The temporal factor probability t was implicitly modeled because t + s + c = 1. Intuitively, β_0_ reflected the temporal factor’s total GM volume of the structures while discounting ICV, β_s_ reflected the response difference between the subcortical and temporal factors, and β_c_ reflected the response difference between the cortical and temporal factors.

Statistical tests of whether total GM volume y varied across factors involved null hypotheses of the form Hβ = 0, where β = [β_0_, β_s_, β_c_, β_i_]^T^, and H is the linear contrast [5]. By specifying different H’s, we were able to compare different pairs of factors. For example, H = [0, 1, 0, 0] tested possible differences between the subcortical and temporal factors, and H = [0, −1, 1, 0] compared the cortical and subcortical factors.

The GLM and statistical tests were repeated using Table S1B (top GM anatomical structures associated with the subcortical factor) and Table S1C (top GM anatomical structures associated with the cortical factor).

**Linear Mixed-Effects Modeling of Longitudinal Cognition Decline Among Aβ+ CN, Aβ+ MCI and AD Dementia Participants.** To analyze variations in cognitive decline rates across atrophy factors, one could first estimate the decline rate for each participant and then model the estimated decline rates using GLM. However, this approach is suboptimal because participants with one or even two time points may have to be discarded because the decline rate cannot be estimated with confidence (e.g., [6]).

Here we considered the linear mixed-effects (LME) model that provides significantly improved exploitation of longitudinal measurements [7] by accounting for both intra-individual measurement correlations and inter-individual variability. Under this framework, the longitudinal cognitive decline rates can be easily compared across atrophy factors for the 188 AD dementia patients, 147 Aβ+ MCI participants, and 43 Aβ+ CN participants.

A single LME model was utilized to examine longitudinal changes in memory (ADNI-Mem [8]) across the atrophy factors in the 43 Aβ+ CN, 147 Aβ+ MCI, and 188 AD dementia patients. The same model was estimated for K = 2, 3 and 4 factors, as well as for executive function (ADNI-EF) and MMSE.

For ease of explanation, we will focus on explaining the LME model for the case of three atrophy factors and ADNI-Mem. Response variable y of the LME model consisted of the 378 (= 43 CN + 147 MCI + 188 AD) participants’ longitudinal ADNI-Mem. Explanatory fixed-effects variables included binary MCI group indicator m, binary AD group indicator d, subcortical factor probability s, cortical factor probability c, interactions between group indicators and factor probabilities (i.e., m·s, m·c, d·s and d·c), time from baseline t, interactions between group indicators and time from baseline (i.e., m·t and d·t), interactions between factor probabilities and time from baseline (i.e., s·t and c·t), and interactions among group indicators, factor probabilities and time from baseline (i.e., m·s·t, m·c·t, d·s·t and d·c·t), while nuisance variables consisted of baseline age x_1_, sex x_2_, education x_3_ and total atrophy x_4_.

The resulting LME model was y = (β_0_ + β_m_·m + β_d_·d + β_s_·s + β_c_·c + β_ms_·m·s + β_mc_·m·c + β_ds_·d·s + β_dc_·d·c + β_1_·x_1_ + β_2_·x_2_ + β_3_·x_3_ + β_4_·x_4_ + b) + (β_t0_ + β_tm_·m + β_td_·d + β_ts_·s + β_tc_·c + β_tms_·m·s + β_tmc_·m·c + β_tds_·d·s + β_tdc_·d·c)·t + ε, where β’s are the regression coefficients, b is the random intercept, and ε is the residual. For the same reasons provided in the previous section, the temporal factor probability and binary CN group indicator were implicitly modeled. Intuitively, Pto reflected the temporal factor’s decline rate at the CN stage, β_t0_ + β_tm_ reflected the temporal factor’s decline rate at the MCI stage, and β_t0_ + β_tm_ + β_ts_ + β_tms_ reflected the subcortical factor’s decline rate at the MCI stage. With this model setup, variations in age, sex, education and total atrophy were controlled for across participants.

Statistical tests were performed in two stages. First, we tested whether ADNI-Mem decline rate accelerated, decelerated or stayed the same across disease stages for each factor. More specifically, for each factor, we first tested whether decline in memory and executive function was significant at the CN stage and then examined possible changes in decline rates from CN to MCI as well as from MCI to AD. For example, to test whether ADNI-Mem decline was significant at the CN stage for the subcortical factor, the null hypothesis was β_t_ + β_ts_ = 0. To test whether the decline rate changed from CN to MCI for the subcortical factor, the null hypothesis was β_mt_ + β_mst_ = 0. Finally, null hypothesis βts + β_dst_−β_mt_−β_mst_ = 0 tested whether the decline accelerated from MCI to AD. The test results for memory and executive function are shown in Figs. 5A2 and 5B2, respectively. Details on hypothesis testing in the LME model can be found in [7].

To foreshadow the results, the hypothesis tests in the previous paragraph hinted at differences in ADNI-Mem decline rates across the factors. Therefore, statistical tests of whether ADNI-mem decline rates varied across factors at each disease stage were performed. More specifically, at each disease stage, we first performed an omnibus statistical test on whether there were differences in memory decline rates across factors and then tested for pairwise differences. Take the MCI stage as an example. Rejecting the null hypothesis β_ts_ + β_tms_ = β_tc_ + β_tmc_ = 0 would imply differences in ADNI-Mem decline rates across the three factors among Aβ+ MCI participants. Rejecting the null hypothesis that β_ts_ + β_tms_ = 0 would suggest that the subcortical factor and temporal factor were associated with different ADNI-Mem decline rates. Rejecting the null hypothesis that β_tc_ + β_tmc_ = 0 would suggest that the cortical factor and temporal factor were associated with different ADNI-Mem decline rates. Finally, rejecting the null hypothesis that β_ts_ + β_tms_ = β_tc_ + β_tmc_ would suggest that the subcortical and cortical factors were associated with different cognitive decline rates.

The results of the above statistical tests are illustrated in Figs. 5A2, 5B2, 7, S6A2, S6B2, S7A2, S7B2, S7C2, S8A2 and S8B2, where (except in Figs. 5A2 and 5B2) the blue dot corresponds to the estimated difference in cognitive decline rate between two “pure factors” after controlling for age, sex, education and total atrophy. For example, when comparing temporal and subcortical factors at the AD dementia stage, the estimated difference in cognitive decline rate is given by β_ts_ + β_tds_. The red bar corresponds to the standard error of this estimation given by std(β_ts_ + β_tds_).

---

**Fig. S1.**
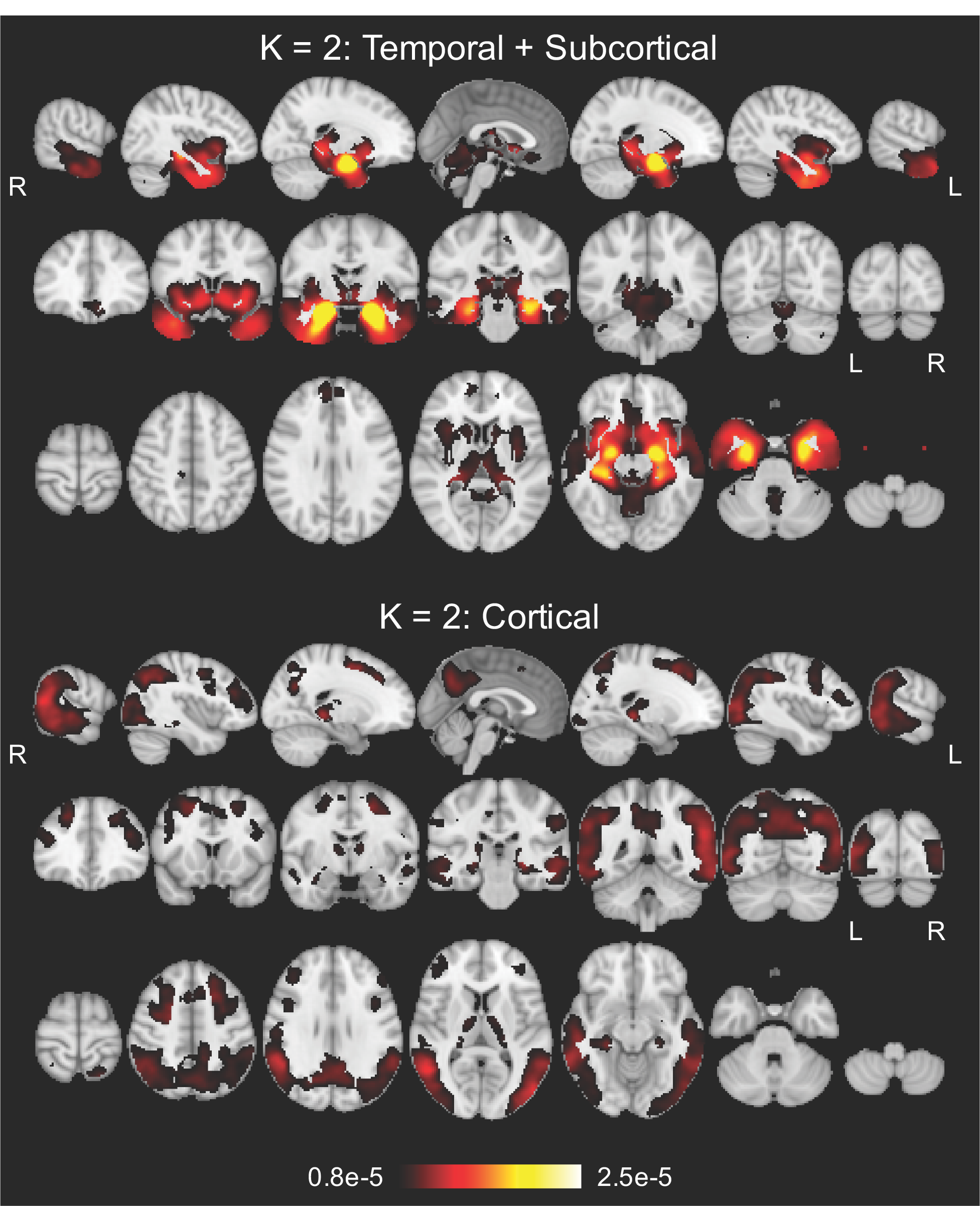
Sagittal, coronal and axial slices of the probabilistic atrophy maps for K = 2, 3 and 4 atrophy factors. Bright color indicates high probability of atrophy at that spatial location for a particular atrophy factor, i.e., Pr(Voxel | Factor).

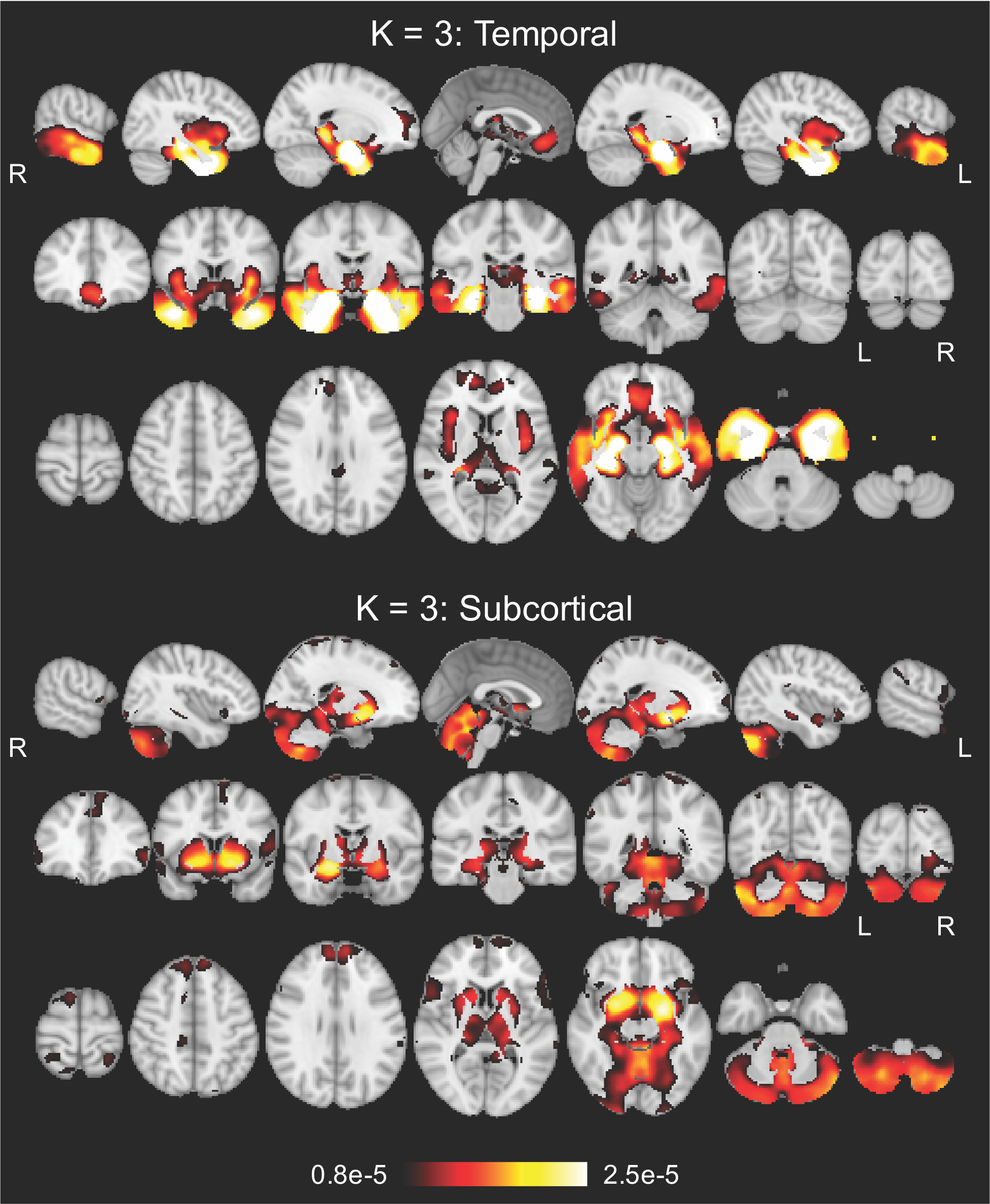

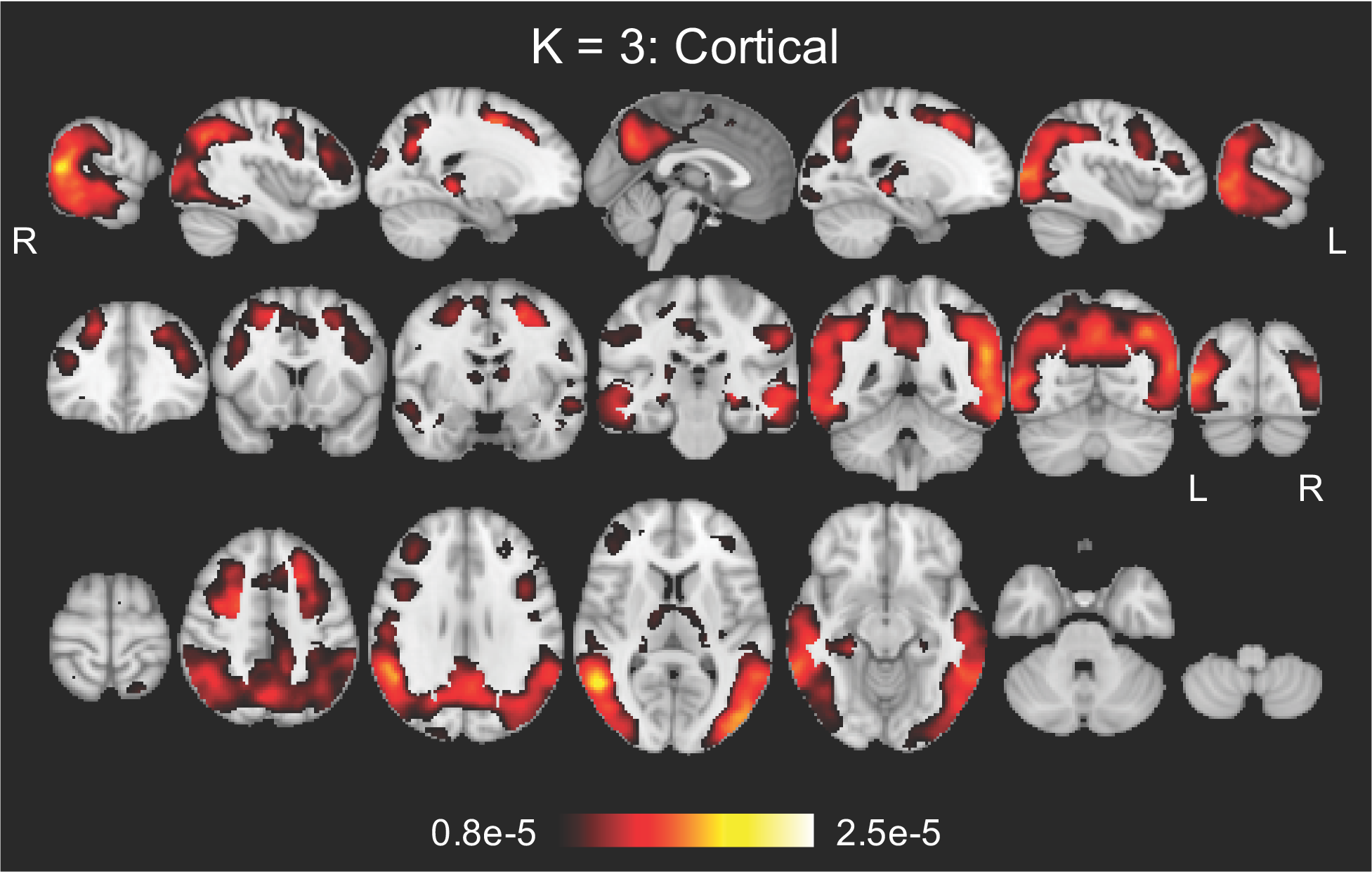

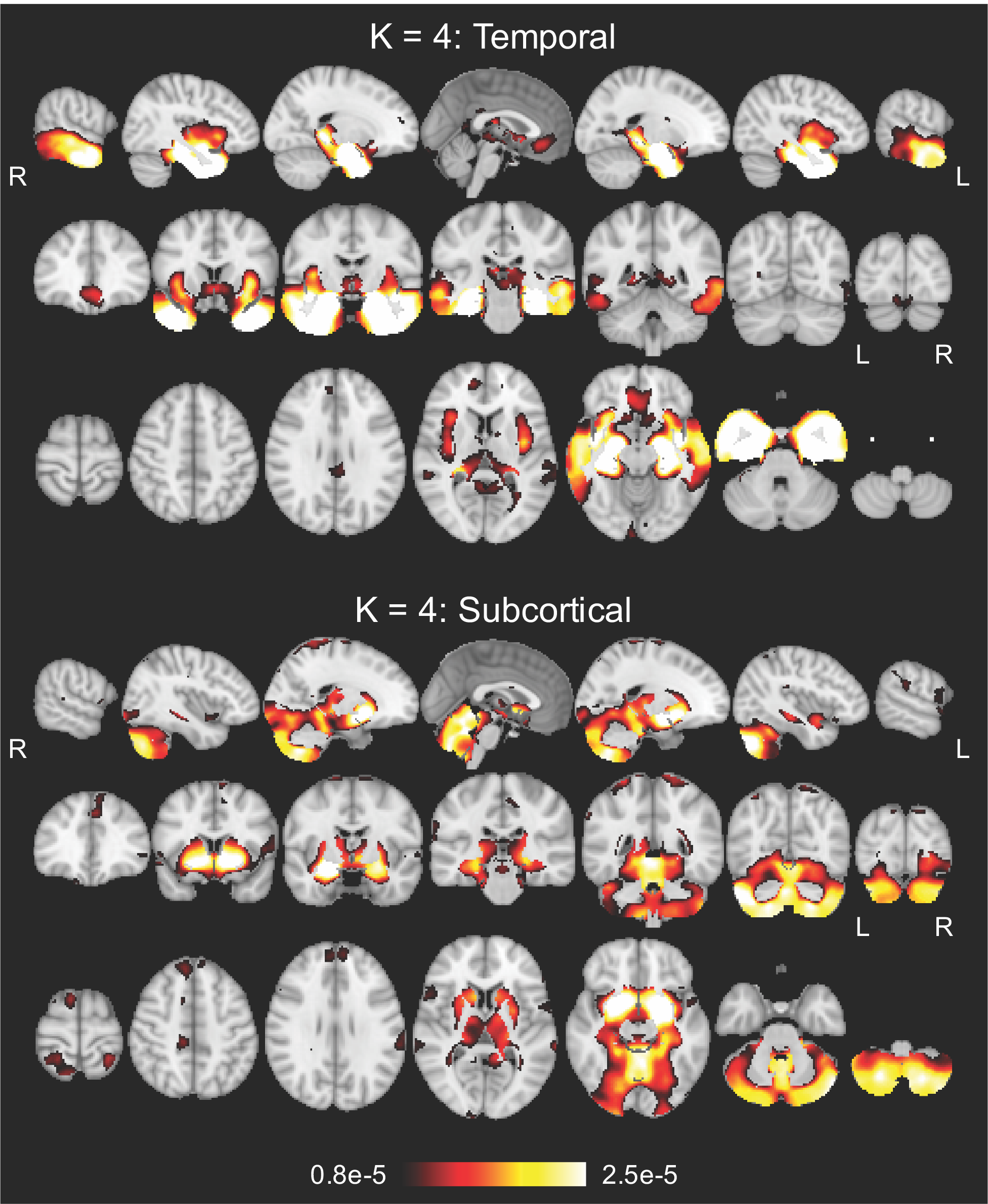

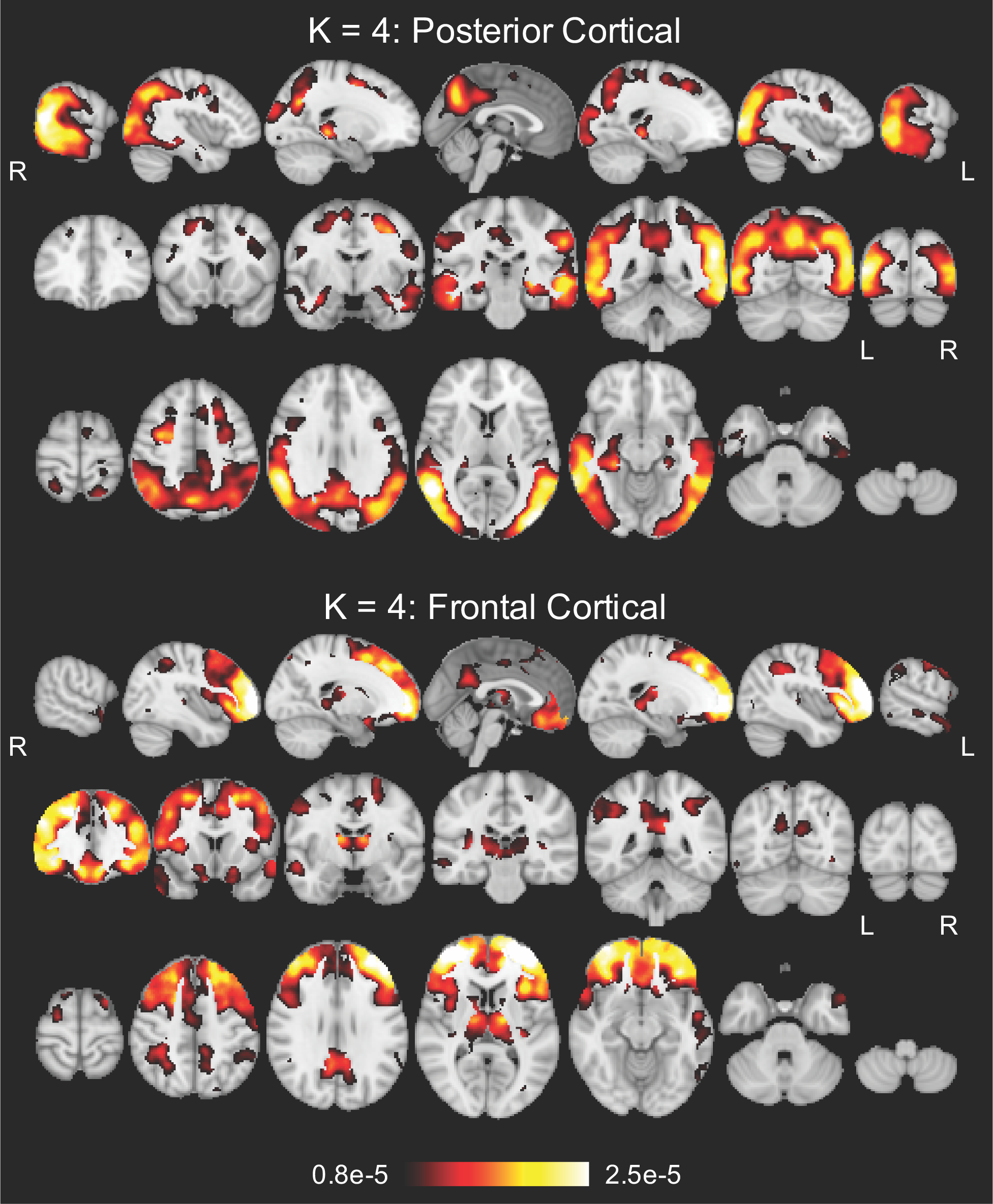

**Fig. S2.**
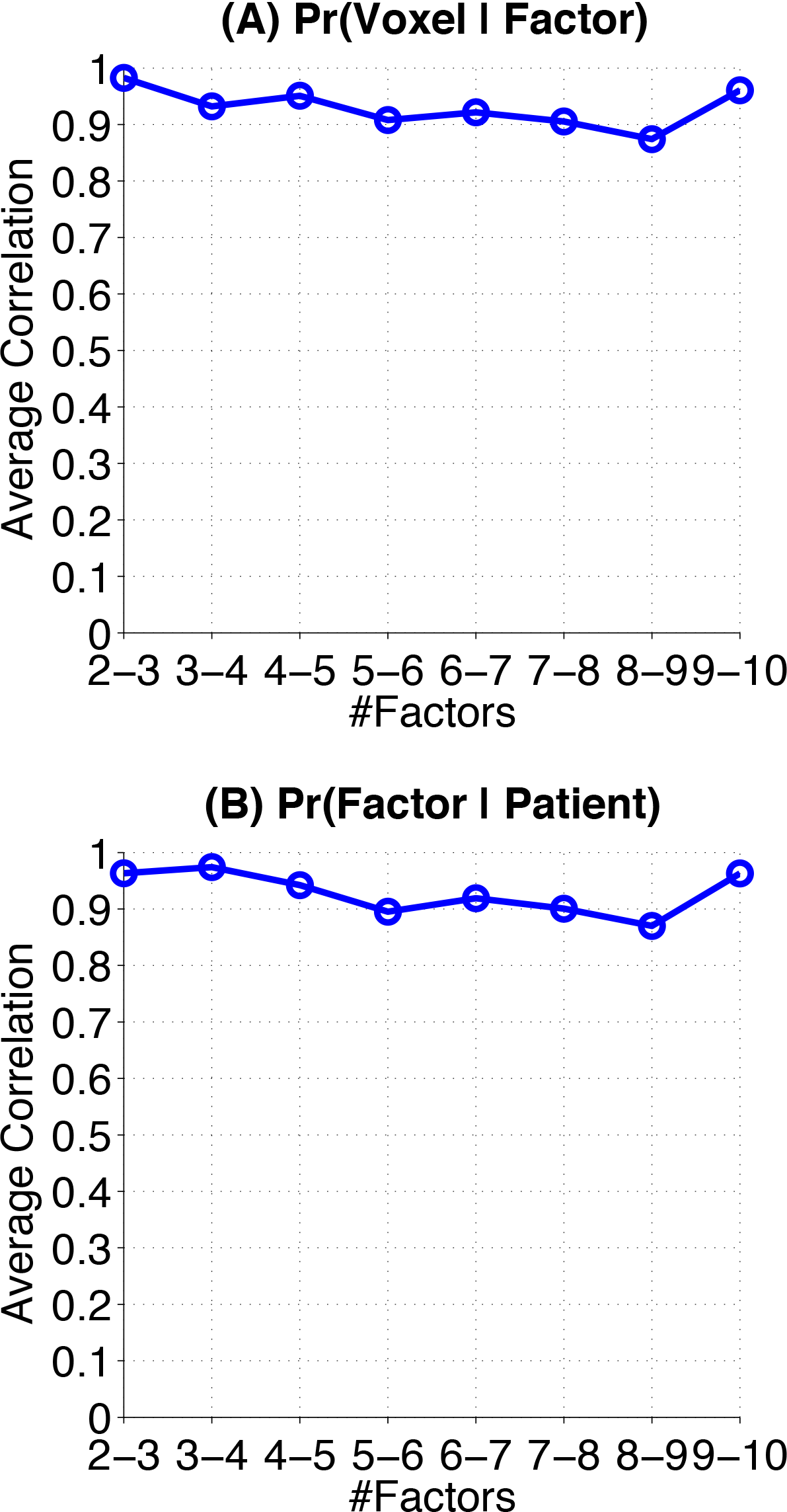
Quantifying the nested hierarchy of latent atrophy factors in terms of (A) atrophy patterns and (B) individual factor compositions. A high correlation value at “K-(K+1)” on the x-axis indicates a high-quality split from the K-factor model to the (K+1)-factor model (see **Supplemental Methods** of **SI**). For example, the close-to-one values at “2-3” in both (A) and (B) suggest that the splits of both the atrophy patterns and individual factor compositions are high-quality from two to three atrophy factors. Overall, the high correlation values from 2 to 10 support a nested hierarchy of latent atrophy factors.

**Fig. S3.**
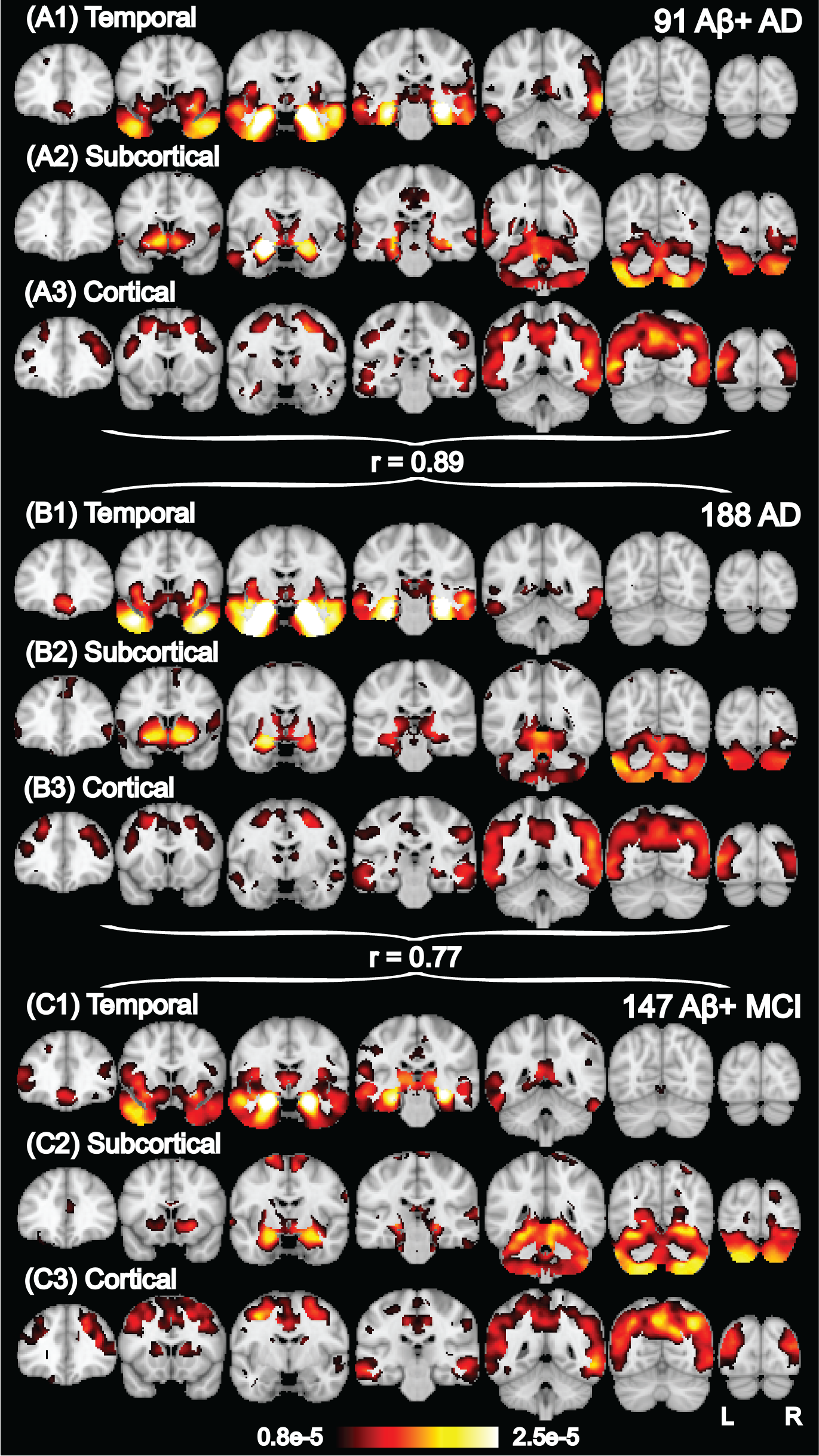
Probabilistic atrophy maps for K = 3 factors estimated with (A) 91 Aβ+AD dementia patients, (B) all 188 AD dementia patients, and (C) 147 Aβ + MCI participants. The three different cohorts yielded highly similar atrophy patterns. Bright color indicates high probability of atrophy at that spatial location for a particular atrophy factor, i.e., Pr(Voxel | Factor).

**Table S1A.** Top anatomical structures associated with the temporal factor (see **Methods**). The temporal factor was associated with significantly greater atrophy in these structures than the subcortical factor (p = 2e-15) and cortical factor (p = 4e-15). There were no differences in atrophy of these structures between the subcortical and cortical factors (p = 0.84). See **Supplemental Methods** of **SI**.

| FreeSurfer Structure Name | Average Probability |
| --- | --- |
| Right-Amygdala | 3.81e-5 |
| Left-Amygdala | 3.59e-5 |
| ctx-rh-entorhinal | 3.03e-5 |
| ctx-lh-entorhinal | 2.87e-5 |
| Right-Hippocampus | 2.86e-5 |
| Left-Hippocampus | 2.51e-5 |
| ctx-rh-parahippocampal | 2.24e-5 |
| ctx-lh-temporalpole | 2.06e-5 |
| ctx-rh-temporalpole | 1.95e-5 |
| ctx-lh-parahippocampal | 1.78e-5 |
| ctx-rh-inferiortemporal | 1.52e-5 |
| ctx-lh-middletemporal | 1.50e-5 |
| ctx-rh-middletemporal | 1.47e-5 |
| ctx-rh-fusiform | 1.40e-5 |
| ctx-lh-inferiortemporal | 1.32e-5 |
| ctx-lh-fusiform | 1.26e-5 |
| ctx-rh-insula | 1.26e-5 |
| ctx-lh-insula | 1.20e-5 |
| ctx-lh-superiortemporal | 1.09e-5 |
| ctx-lh-rostralanteriorcingulate | 1.03e-5 |
| ctx-rh-superiortemporal | 9.82e-6 |
| ctx-rh-medialorbitofrontal | 8.39e-6 |
| ctx-rh-rostralanteriorcingulate | 7.77e-6 |
| ctx-rh-lateralorbitofrontal | 7.71e-6 |
| ctx-lh-medialorbitofrontal | 7.71e-6 |
| ctx-rh-transversetemporal | 7.13e-6 |
| ctx-lh-lateralorbitofrontal | 6.92e-6 |
| Right-VentralDC | 5.95e-6 |
| ctx-lh-caudalanteriorcingulate | 3.71e-6 |

**Table S1B.** Top anatomical structures associated with the subcortical factor (see **Methods**). The subcortical factor was associated with significantly greater atrophy in these structures than the temporal factor (p = 1e-5) and cortical factor (p = 2e-12). The temporal factor had more atrophy in these structures than the cortical factor (p = 0.01). See **Supplemental Methods** of **SI**.

| FreeSurfer Structure Name | Average Probability |
| --- | --- |
| Right-Accumbens-area | 1.85e-5 |
| Left-Accumbens-area | 1.75e-5 |
| Right-Putamen | 1.31e-5 |
| Left-Cerebellum-Cortex | 1.16e-5 |
| Left-Putamen | 1.13e-5 |
| Right-Cerebellum-Cortex | 1.10e-5 |
| Left-Thalamus-Proper | 8.82e-6 |
| Right-Thalamus-Proper | 7.99e-6 |
| Right-Caudate | 7.62e-6 |
| ctx-lh-lingual | 7.58e-6 |
| Left-Caudate | 7.50e-6 |
| ctx-rh-lingual | 7.16e-6 |
| ctx-lh-parstriangularis | 7.10e-6 |
| ctx-rh-parstriangularis | 6.52e-6 |
| ctx-rh-parsopercularis | 6.25e-6 |
| ctx-rh-superiorfrontal | 5.81e-6 |
| ctx-rh-parsorbitalis | 5.57e-6 |
| Left-VentralDC | 5.46e-6 |
| ctx-lh-parsorbitalis | 5.26e-6 |
| ctx-lh-superiorfrontal | 5.01e-6 |
| ctx-lh-frontalpole | 4.31e-6 |
| ctx-rh-frontalpole | 3.57e-6 |
| Brain-Stem | 3.36e-6 |
| Right-Pallidum | 2.55e-6 |
| Left-Pallidum | 2.22e-6 |

**Table S1C.** Top anatomical structures associated with the cortical factor (see **Methods**). The cortical factor was associated with significantly greater atrophy in these structures than the temporal factor (p = 7e-6) and subcortical factor (p = 4e-7). There were no differences in atrophy of these structures between the temporal and subcortical factors (p = 0.62). See **Supplemental Methods** of **SI**.

| FreeSurfer Structure Name | Average Probability |
| --- | --- |
| ctx-lh-bankssts | 1.15e-5 |
| ctx-rh-inferiorparietal | 1.10e-5 |
| ctx-lh-precuneus | 1.00e-5 |
| ctx-rh-bankssts | 9.92e-6 |
| ctx-rh-precuneus | 9.07e-6 |
| ctx-lh-inferiorparietal | 8.94e-6 |
| ctx-lh-caudalmiddlefrontal | 8.47e-6 |
| ctx-rh-caudalmiddlefrontal | 8.37e-6 |
| ctx-rh-lateraloccipital | 8.22e-6 |
| ctx-lh-supramarginal | 7.99e-6 |
| ctx-lh-lateraloccipital | 7.64e-6 |
| ctx-rh-isthmuscingulate | 7.32e-6 |
| ctx-rh-cuneus | 7.16e-6 |
| ctx-lh-isthmuscingulate | 7.11e-6 |
| ctx-lh-superiorparietal | 6.89e-6 |
| ctx-rh-supramarginal | 6.74e-6 |
| ctx-lh-paracentral | 6.53e-6 |
| ctx-lh-cuneus | 6.47e-6 |
| ctx-lh-transversetemporal | 6.29e-6 |
| ctx-rh-posteriorcingulate | 6.29e-6 |
| ctx-lh-parsopercularis | 6.05e-6 |
| ctx-lh-posteriorcingulate | 5.87e-6 |
| ctx-lh-rostralmiddlefrontal | 5.69e-6 |
| ctx-rh-precentral | 5.69e-6 |
| ctx-rh-superiorparietal | 5.57e-6 |
| ctx-rh-rostralmiddlefrontal | 5.41e-6 |
| ctx-lh-precentral | 5.33e-6 |
| ctx-lh-pericalcarine | 5.29e-6 |
| ctx-lh-postcentral | 5.27e-6 |
| ctx-rh-pericalcarine | 4.94e-6 |
| ctx-rh-postcentral | 4.73e-6 |
| ctx-rh-paracentral | 4.68e-6 |
| ctx-rh-caudalanteriorcingulate | 3.83e-6 |

**Fig. S4.**
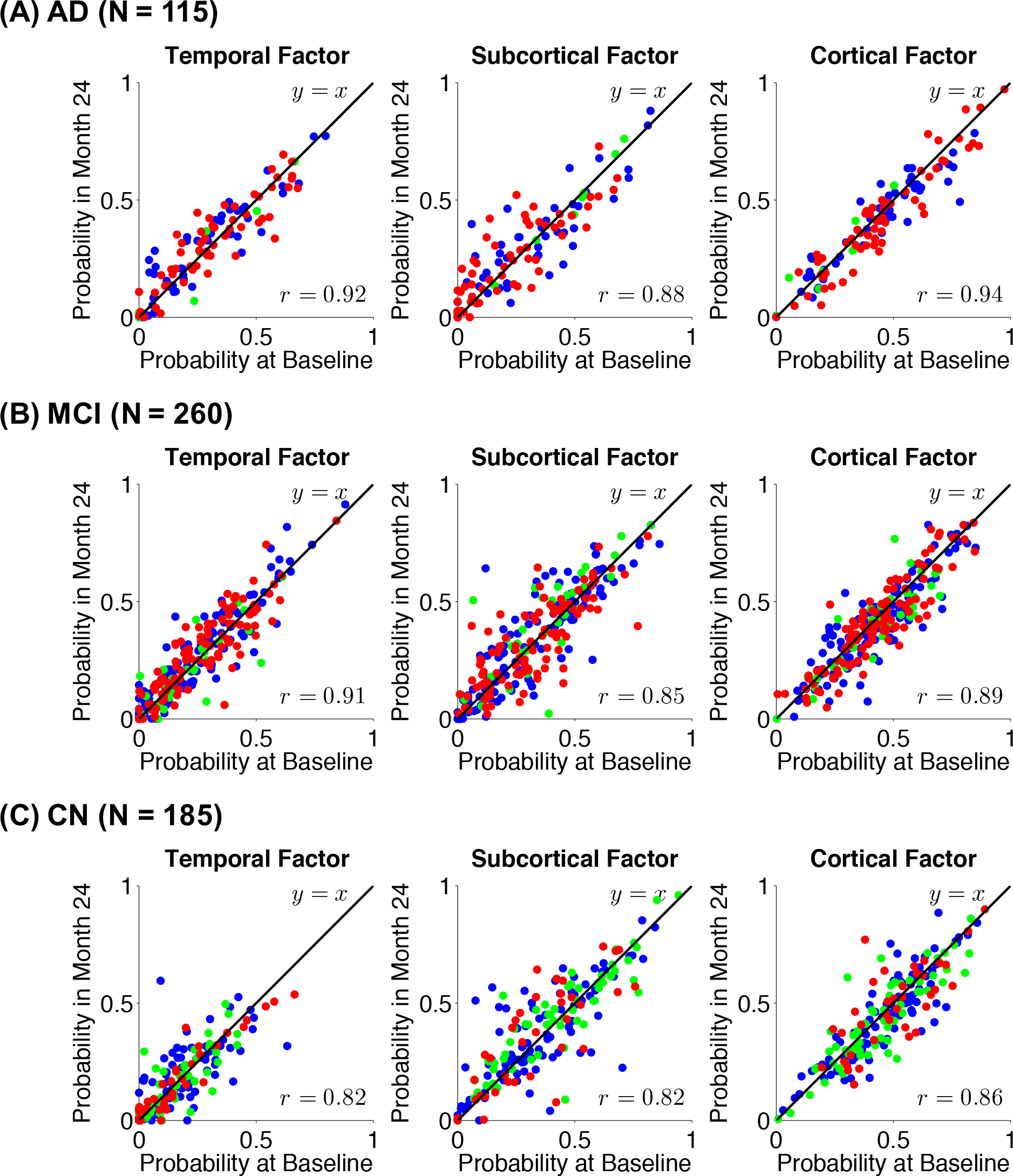
Stability of factor compositions over two years for (A) 115 AD dementia patients, 260 MCI participants, and (C) 185 CN participants. Each participant corresponds to a dot, whose color indicates amyloid status: red for Aβ+, green for Aβ-, and blue for unknown. For each atrophy factor (plot), x-axis and y-axis represent, respectively, the probabilities of factor at baseline and two years after baseline. In the ideal case where factor probability estimations remain exactly the same after two years, one would expect a y = x linear fit as well as a r = 1 correlation. In our case, the linear fits were close to y = x with r > 0.82 for all three atrophy factors for all clinical groups, suggesting that the factor compositions were stable despite disease progression.

**Fig. S5A.**
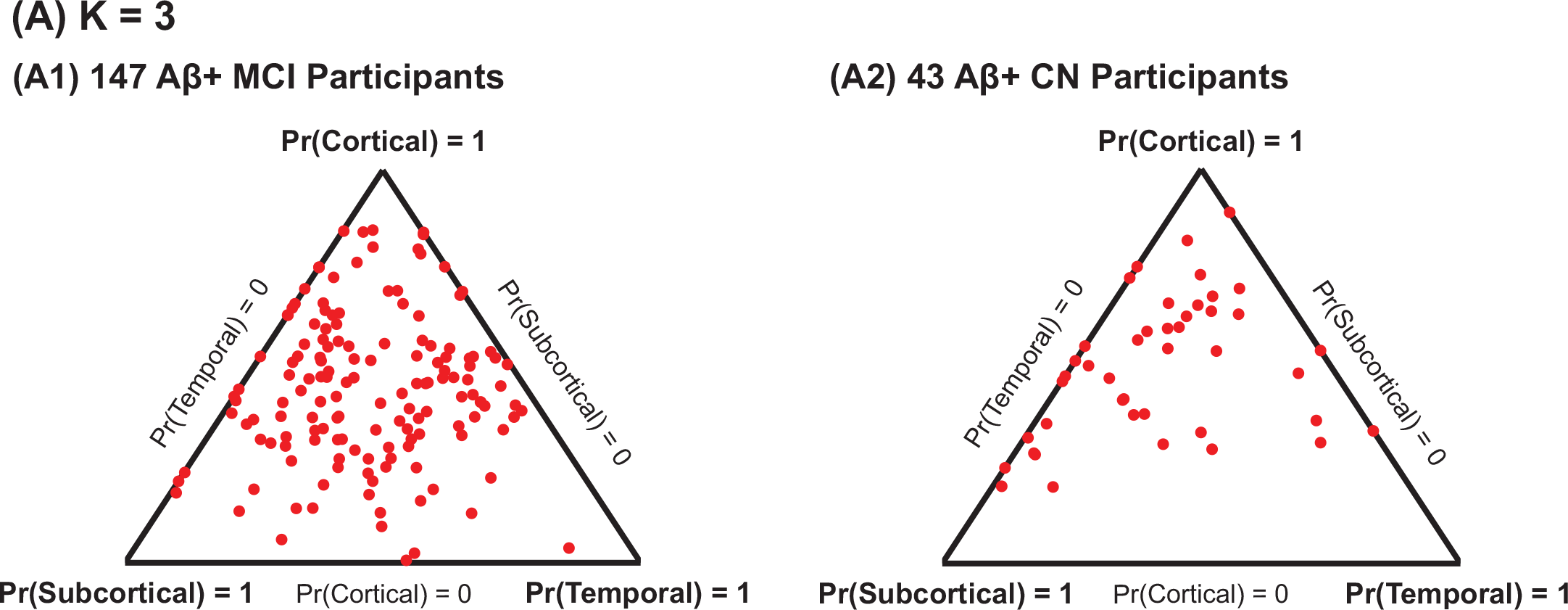
Factor compositions of (1) 147 Aβ+MCI participants and (2) 43 Aβ+CN participants for K = 3 factors. Each participant corresponds to a dot, whose location (in barycentric coordinates) represents the factor composition. Corners of the triangle represent “pure factors” closer distance to the respective corners indicates higher probabilities for the respective factors. Most dots are far from the corners, suggesting that most participants expressed multiple factors.

**Fig. S5B.**
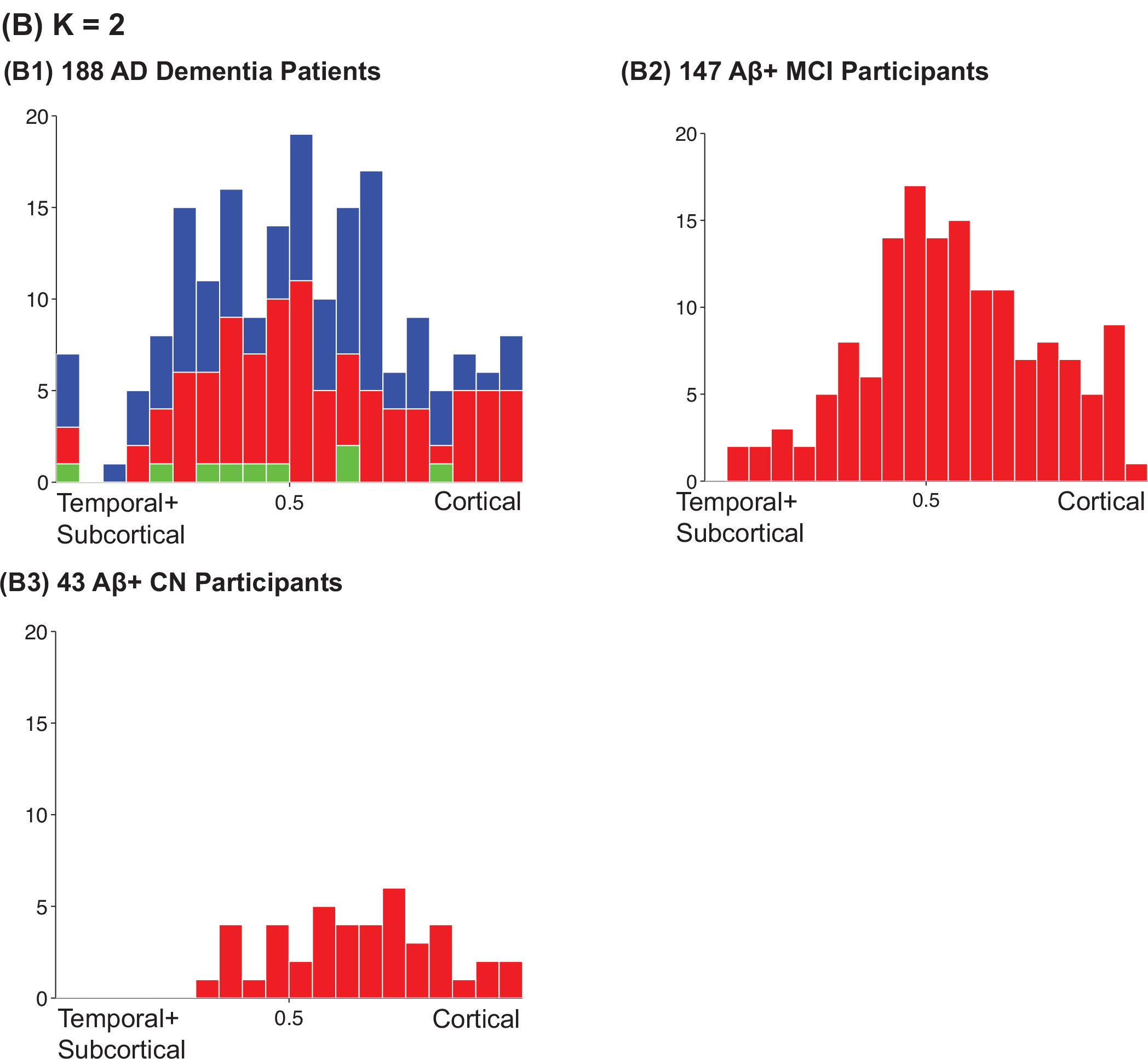
Factor compositions of (1) 188 AD dementia patients, (2) 147 Aβ+ MCI participants, and (3) 43 Aβ+ CN participants for K = 2 factors. Histograms were created with participants’ cortical factor probability (x-axis). Therefore the left (or right) extreme corresponds to the pure temporal+subcortical (or cortical) factor. In addition, colors in (1) indicate amyloid status: red for Aβ+, green for Aβ−, and blue for unknown. The majority of the population lies around the center, suggesting that most participants expressed both atrophy factors.

**Fig. S5C.**
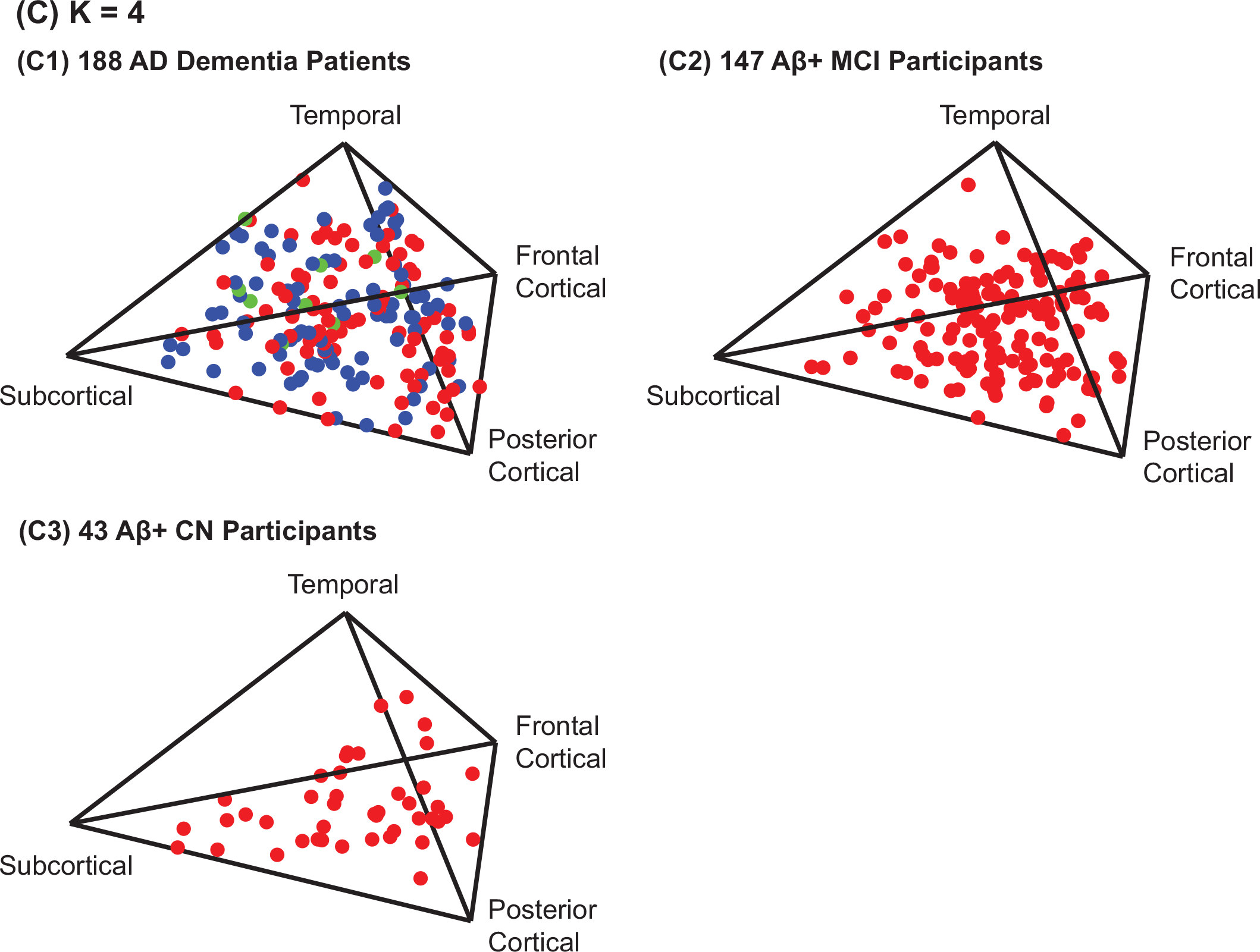
Factor compositions of (1) 188 AD dementia patients, (2) 147 Aβ+MCI participants, and (3) 43 Aβ+ CN participants for K = 4 factors. Each participant corresponds to a dot, whose location represents the factor composition. Tetrahedron corners represent “pure factors”; closer distance to a corner corresponds to higher probability for the corresponding factor. Color in (1) indicates amyloid status: red for Aβ+, green for Aβ−, and blue for unknown. Most dots are far from the corners, suggesting that most participants expressed multiple factors.

**Table S2.**
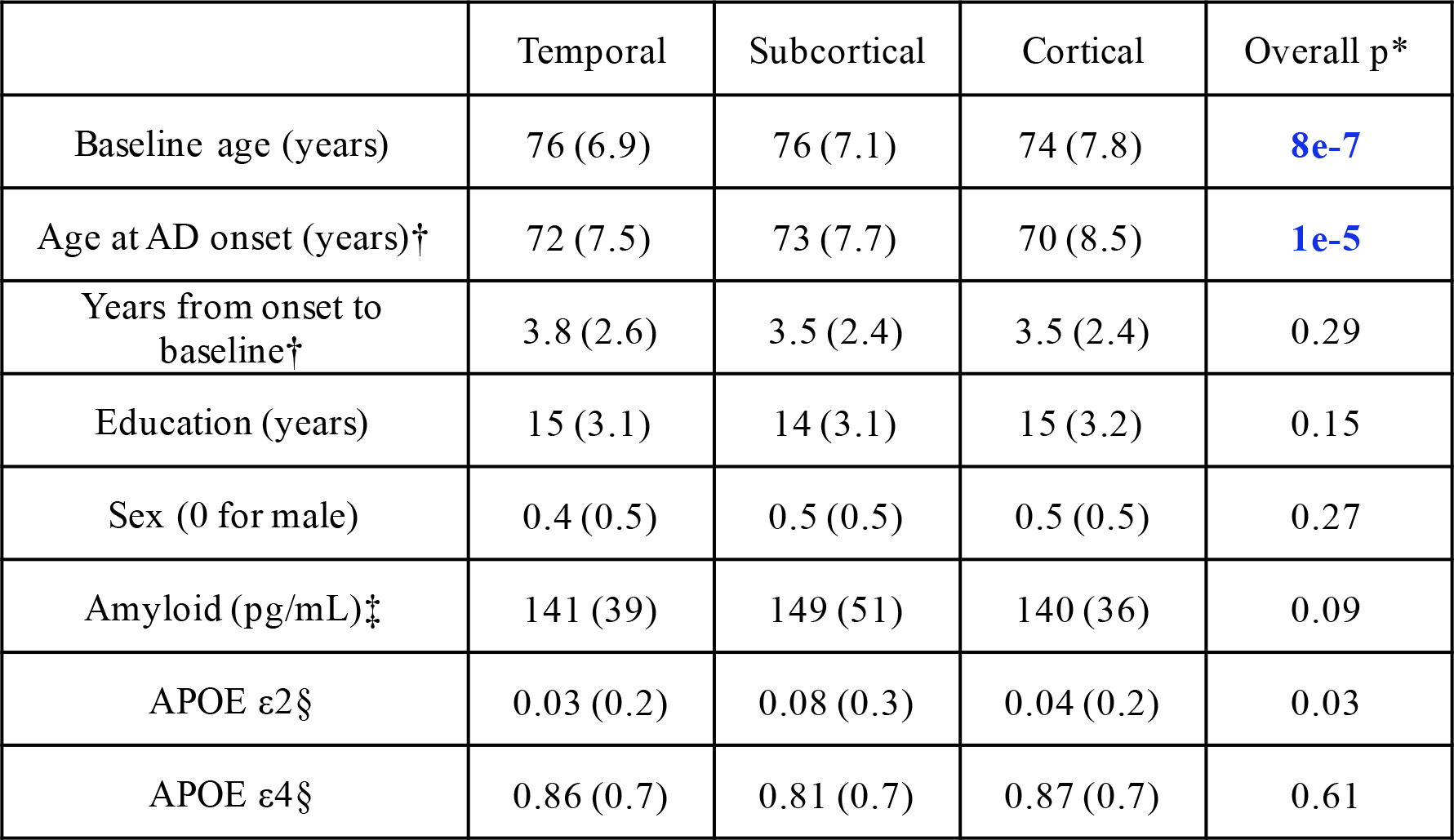
Characteristics of 188 AD dementia patients by factor. Data are weighted averages (weighted standard deviation) with weights corresponding to factor probabilities. Highlighted p values (blue) are characteristics significantly different across factors. * Computed by linear hypothesis test on GLM or likelihood ratio test on logistic regression for sex (see **Methods**) † Only available for 182 patients. ‡Only available for 100 patients. § The original counts were 0, 1 or 2.

|  | Temporal | Subcortical | Cortical | Overall p* |
| --- | --- | --- | --- | --- |
| Baseline age (years) | 76 (6.9) | 76 (7.1) | 74 (7.8) | **8e-7** |
| Age at AD onset (years)† | 72 (7.5) | 73 (7.7) | 70 (8.5) | **1e-5** |
| Years from onset to baseline† | 3.8 (2.6) | 3.5 (2.4) | 3.5 (2.4) | 0.29 |
| Education (years) | 15 (3.1) | 14 (3.1) | 15 (3.2) | 0.15 |
| Sex (0 for male) | 0.4 (0.5) | 0.5 (0.5) | 0.5 (0.5) | 0.27 |
| Amyloid (pg/mL)‡ | 141 (39) | 149 (51) | 140 (36) | 0.09 |
| APOE ε2§ | 0.03 (0.2) | 0.08 (0.3) | 0.04 (0.2) | 0.03 |
| APOE ε4§ | 0.86 (0.7) | 0.81 (0.7) | 0.87 (0.7) | 0.61 |

**Fig. S6.**
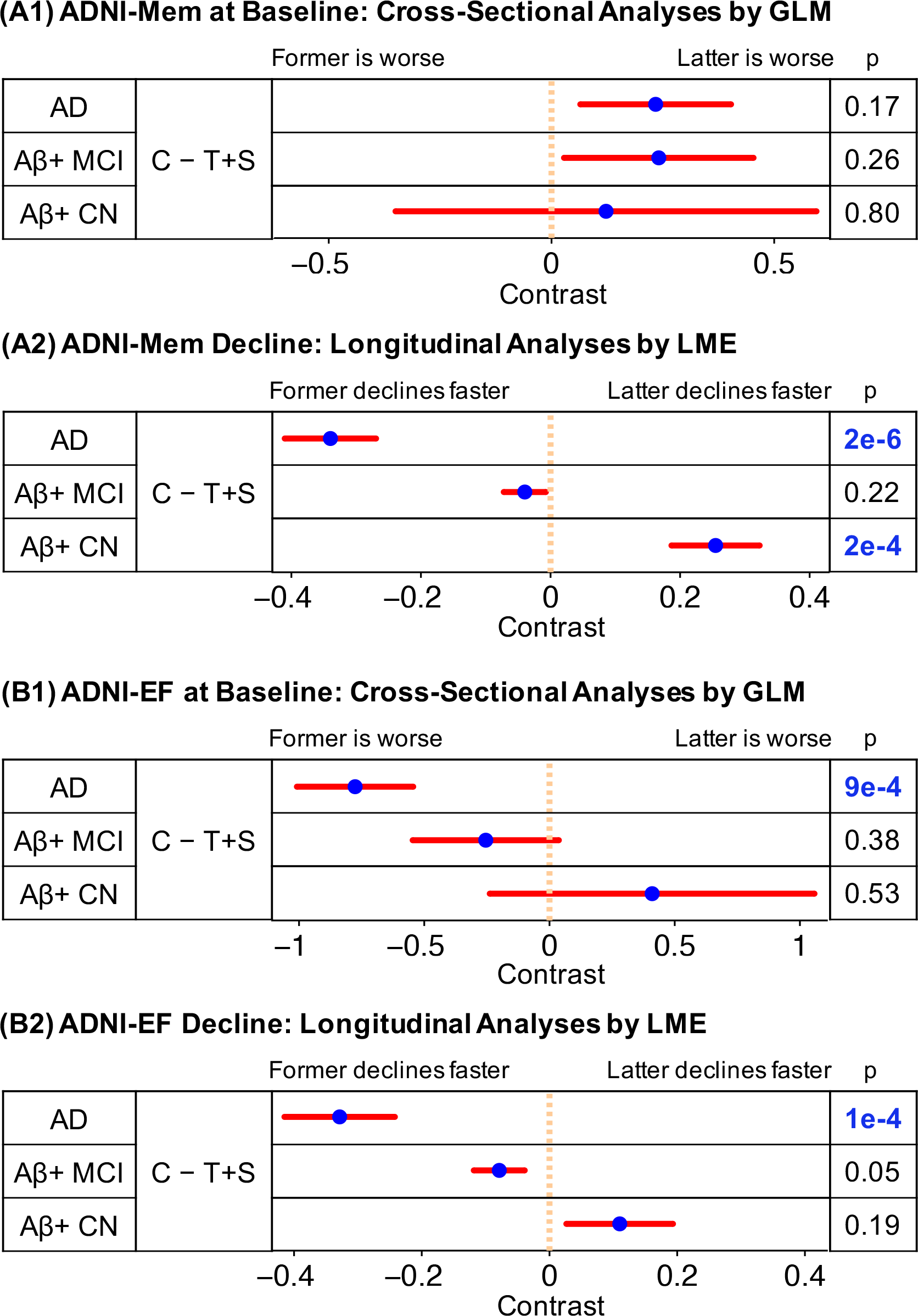
Comparisons of (1) cross-sectional baseline and (2) longitudinal decline rates of (A) memory and (B) executive function between K = 2 factors. Comparisons remaining significant after FDR control (q = 0.05) are highlighted in blue. Blue dots are estimated differences between “pure atrophy factors”, and red bars show the standard errors (see **Methods** and **Supplemental Methods** of **SI**).

**Fig. S7A.**
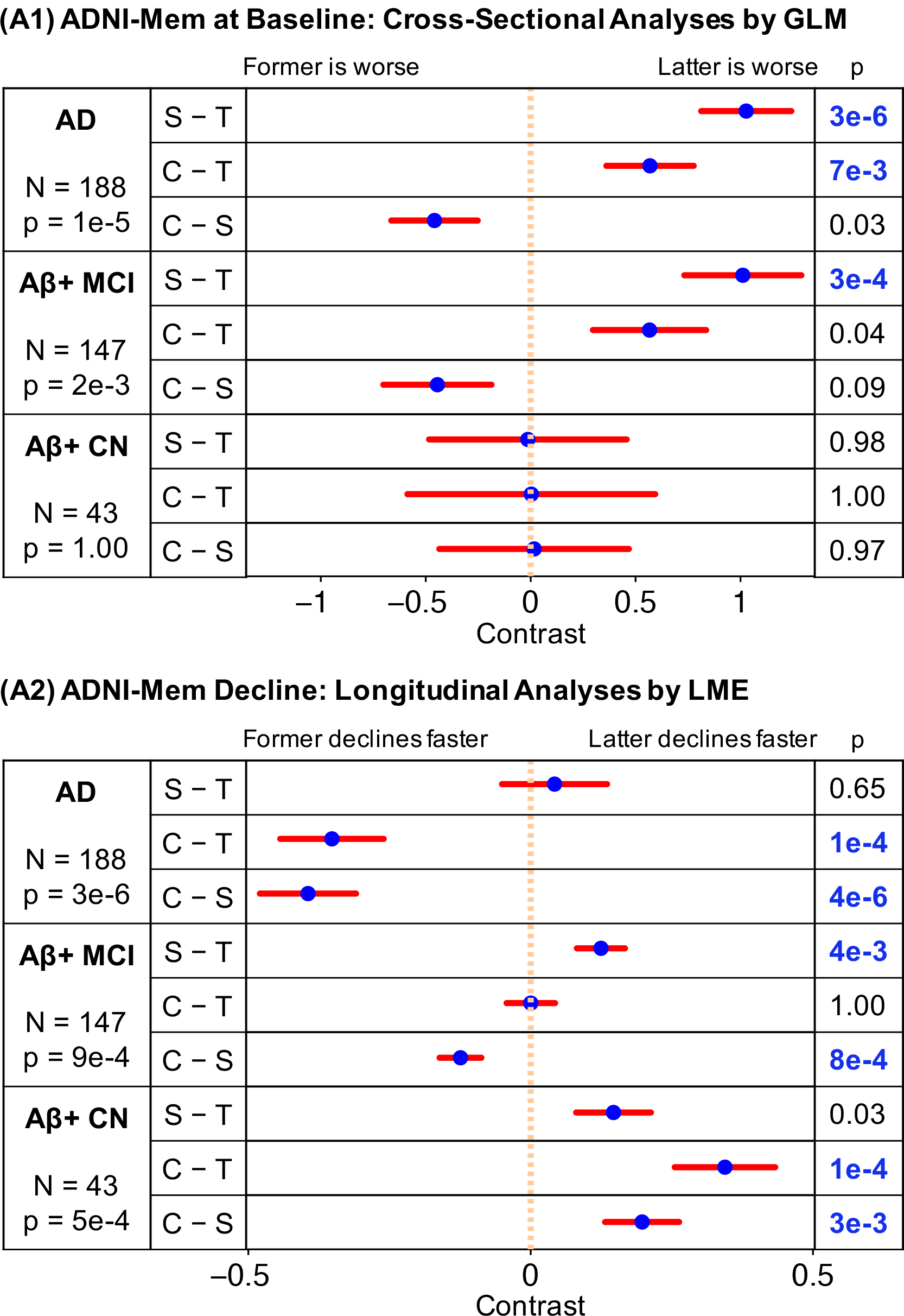
Comparisons of (1) cross-sectional baseline and (2) longitudinal decline rates of memory among K = 3 factors. Comparisons remaining significant after FDR control (q = 0.05) are highlighted in blue. Blue dots are estimated differences between “pure atrophy factors”, and red bars show the standard errors (see **Methods** and **Supplemental Methods** of **SI**).

**Fig. S7B.**
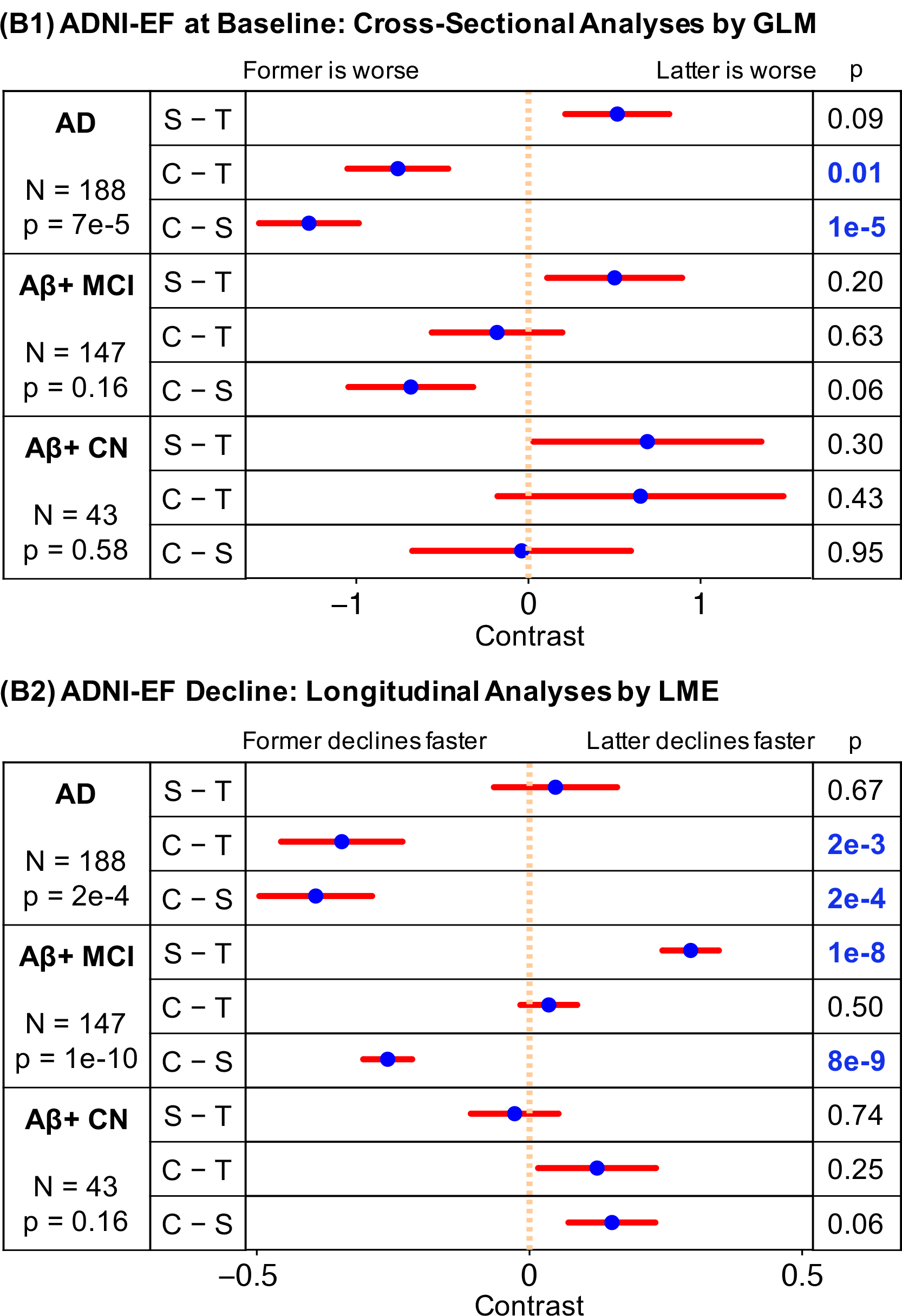
Comparisons of (1) cross-sectional baseline and (2) longitudinal decline rates of executive function among K = 3 factors. Comparisons remaining significant after FDR control(q= 0.05) are highlighted in blue. Blue dots are estimated differences between “pure atrophy factors”, and red bars show the standard errors (see **Methods** and **Supplemental Methods** of **SI**).

**Fig. S7C.**
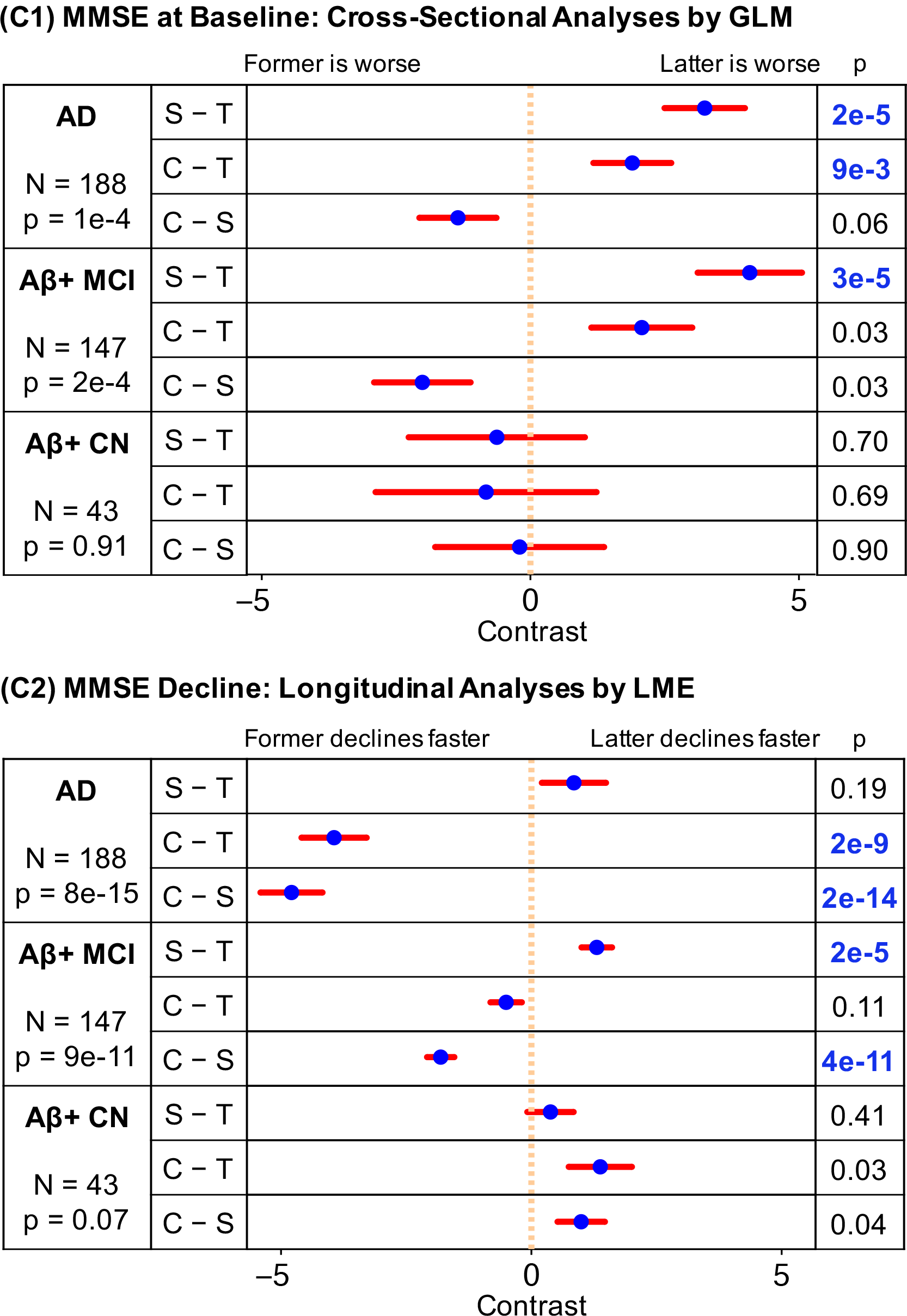
Comparisons of (1) cross-sectional baseline and (2) longitudinal decline rates of MMSE among K = 3 factors. Comparisons remaining significant after FDR control (q = 0.05) are highlighted in blue. Blue dots are estimated differences between “pure atrophy factors”, and red bars show the standard errors (see **Methods** and **Supplemental Methods** of **SI**).

**Fig. S8A.**
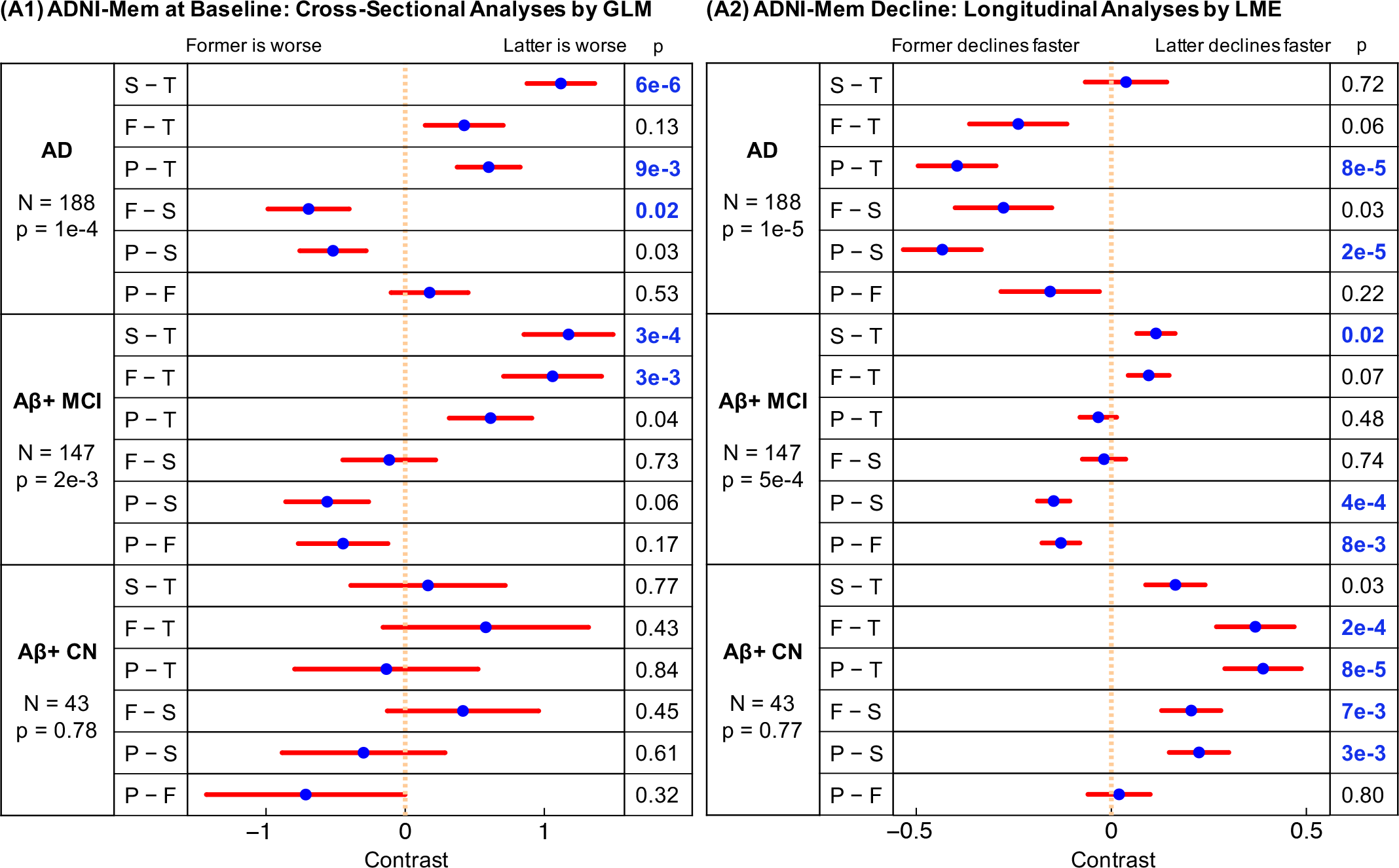
Comparisons of (1) cross-sectional baseline and (2) longitudinal decline rates of MMSE among K = 3 factors. Comparisons remaining significant after FDR control (q = 0.05) are highlighted in blue. Blue dots are estimated differences between “pure atrophy factors”, and red bars show the standard errors (see **Methods** and **Supplemental Methods** of **SI**).

**Fig. S8B.**
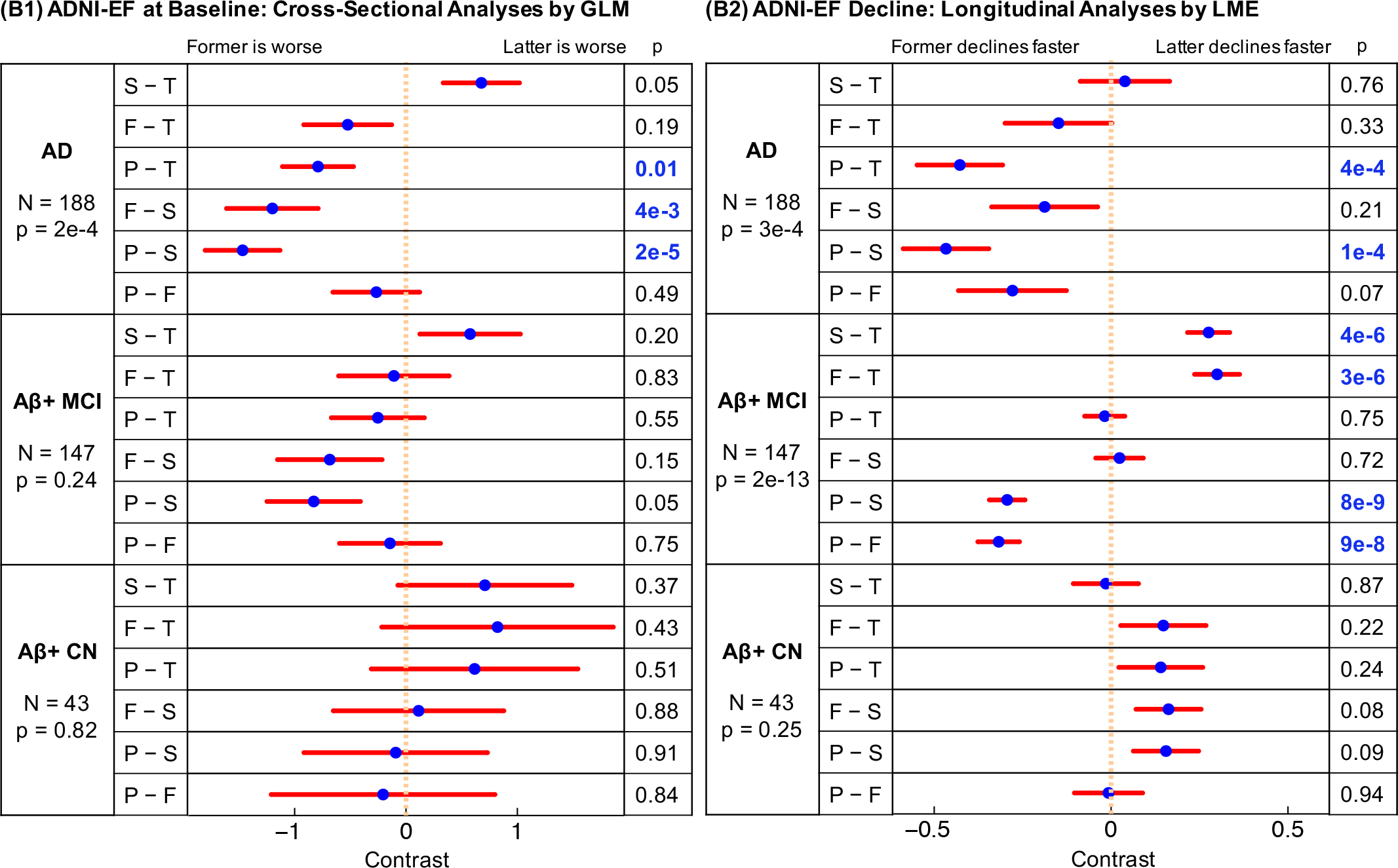
Comparisons of (1) cross-sectional baseline and (2) longitudinal decline rates of MMSE among K = 3 factors. Comparisons remaining significant after FDR control (q = 0.05) are highlighted in blue. Blue dots are estimated differences between “pure atrophy factors”, and red bars show the standard errors (see **Methods** and **Supplemental Methods** of **SI**).

## Complete List of ADNI Investigators and Participating Institutions

### ADNI Investigators

The following list is also available online at http://adni.loni.usc.edu/wp-content/uploads/how_to_apply/ADNI_Acknowledgement_List.pdf.

#### Part A: Leadership and Infrastructure

Principal Investigator
Michael W. Weiner, MD UC San Francisco

ADCS PI and Director of Coordinating Center Clinical Core
Paul Aisen, MD University of Southern California

Executive Committee
Michael Weiner, MD UC San Francisco
Paul Aisen, MD University of Southern California
Ronald Petersen, MD, PhD Mayo Clinic, Rochester
Clifford R. Jack, Jr., MD Mayo Clinic, Rochester
William Jagust, MD UC Berkeley
John Q. Trojanowki, MD, PhD U Pennsylvania
Arthur W. Toga, PhD USC
Laurel Beckett, PhD UC Davis
Robert C. Green, MD, MPH Brigham and Women’s Hospital/Harvard Medical School
Andrew J. Saykin, PsyD Indiana University
John Morris, MD Washington University St. Louis
Leslie M. Shaw University of Pennsylvania

ADNI External Advisory Board (ESAB)
Zaven Khachaturian, PhD Prevent Alzheimer’s Disease 2020 (Chair)
Greg Sorensen, MD Siemens
Maria Carrillo, PhD Alzheimer’s Association
Lew Kuller, MD University of Pittsburgh
Marc Raichle, MD Washington University St. Louis
Steven Paul, MD Cornell University
Peter Davies, MD Albert Einstein College of Medicine of Yeshiva University
Howard Fillit, MD AD Drug Discovery Foundation
Franz Hefti, PhD Acumen Pharmaceuticals
David Holtzman, MD Washington University St. Louis
M. Marcel Mesulam, MD Northwestern University
William Potter, MD National Institute of Mental Health
Peter Snyder, PhD Brown University

ADNI 2 Private Partner Scientific Board (PPSB)
Adam Schwartz, MD Eli Lilly (Chair)

Data and Publications Committee
Robert C. Green, MD, MPH BWH/HMS (Chair)

Resource Allocation Review Committee
Tom Montine, MD, PhD University of Washington (Chair)

Clinical Core Leaders
Ronald Petersen, MD, PhD Mayo Clinic, Rochester (Core PI)
Paul Aisen, MD University of Southern California

Clinical Informatics and Operations
Ronald G. Thomas, PhD UC San Diego
Michael Donohue, PhD UC San Diego
Sarah Walter, MSc UC San Diego
Devon Gessert UC San Diego
Tamie Sather, MA UC San Diego
Gus Jiminez, MBS UC San Diego
Archana B. Balasubramanian, PhD UC San Diego
Jennifer Mason, MPH UC San Diego
Iris Sim UC San Diego

Biostatistics Core Leaders and Key Personnel
Laurel Beckett, PhD UC Davis (Core PI)
Danielle Harvey, PhD UC Davis
Michael Donohue, PhD UC San Diego

MRI Core Leaders and Key Personnel
Clifford R. Jack, Jr., MD Mayo Clinic, Rochester (Core PI)
Matthew Bernstein, PhD Mayo Clinic, Rochester
Nick Fox, MD University of London
Paul Thompson, PhD UCLA School of Medicine
Norbert Schuff, PhD UCSF MRI
Charles DeCArli, MD UC Davis
Bret Borowski, RT Mayo Clinic
Jeff Gunter, PhD Mayo Clinic
Matt Senjem, MS Mayo Clinic
Prashanthi Vemuri, PhD Mayo Clinic
David Jones, MD Mayo Clinic
Kejal Kantarci Mayo Clinic
Chad Ward Mayo Clinic

PET Core Leaders and Key Personnel
William Jagust, MD UC Berkeley (Core PI)
Robert A. Koeppe, PhD University of Michigan
Norm Foster, MD University of Utah
Eric M. Reiman, MD Banner Alzheimer’s Institute
Kewei Chen, PhD Banner Alzheimer’s Institute
Chet Mathis, MD University of Pittsburgh
Susan Landau, PhD UC Berkeley

Neuropathology Core Leaders
John C. Morris, MD Washington University St. Louis
Nigel J. Cairns, PhD, FRCPath Washington University St. Louis
Erin Franklin, MS, CCRP Washington University St. Louis
Lisa Taylor-Reinwald, BA, HTL (ASCP)-Past Investigator Washington University St. Louis

Biomarkers Core Leaders and Key Personnel
Leslie M. Shaw, PhD UPenn School of Medicine
John Q. Trojanowki, MD, PhD UPenn School of Medicine
Virginia Lee, PhD, MBA UPenn School of Medicine
Magdalena Korecka, PhD UPenn School of Medicine
Michal Figurski, PhD UPenn School of Medicine

Informatics Core Leaders and Key Personnel
Arthur W. Toga, PhD USC (Core PI)
Karen Crawford USC
Scott Neu, PhD USC

Genetics Core Leaders and Key Personnel
Andrew J. Saykin, PsyD Indiana University
Tatiana M. Foroud, PhD Indiana University
Steven Potkin, MD UC UC Irvine
Li Shen, PhD Indiana University
Kelley Faber, MS, CCRC Indiana University
Sungeun Kim, PhD Indiana University
Kwangsik Nho, PhD Indiana University

Initial Concept Planning & Development
Michael W. Weiner, MD UC San Francisco
Lean Thal, MD UC San Diego
Zaven Khachaturian, PhD Prevent Alzheimer’s Disease 2020

Early Project Proposal Development
Early Project Proposal Development
Leon Thal, MD UC San Diego
Neil Buckholtz National Institute on Aging
Michael W. Weiner, MD UC San Francisco
Peter J. Snyder, PhD Brown University
William Potter, MD National Institute of Mental Health
Steven Paul, MD Cornell University
Marilyn Albert, PhD Johns Hopkins University
Richard Frank, MD, PhD Richard Frank Consulting
Zaven Khachaturian, PhD Prevent Alzheimer’s Disease 2020

NIA
John Hsiao, MD National Institute on Aging

#### Part B: Investigators By Site

Oregon Health & Science University:
Jeffrey Kaye, MD
Joseph Quinn, MD
Lisa Silbert, MD
Betty Lind, BS
Raina Carter, BA-Past Investigator
Sara Dolen, BS-Past Investigator

University of Southern California:
Lon S. Schneider, MD
Sonia Pawluczyk, MD
Mauricio Becerra, BS
Liberty Teodoro, RN
Bryan M. Spann, DO, PhD-Past Investigator

University of California-San Diego:
James Brewer, MD, PhD
Helen Vanderswag, RN
Adam Fleisher, MD-Past Investigator

University of Michigan:
Judith L. Heidebrink, MD, MS
Joanne L. Lord, LPN, BA, CCRC-Past
Investigator

Mayo Clinic, Rochester:
Ronald Petersen, MD, PhD
Sara S. Mason, RN
Colleen S. Albers, RN
David Knopman, MD
Kris Johnson, RN-Past Investigator

Baylor College of Medicine:
Rachelle S. Doody, MD, PhD
Javier Villanueva-Meyer, MD
Valory Pavlik, PhD
Victoria Shibley, MS
Munir Chowdhury, MBBS, MS-Past Investigator
Susan Rountree, MD-Past Investigator
Mimi Dang, MD-Past Investigator

Columbia University Medical Center:
Yaakov Stern, PhD
Lawrence S. Honig, MD, PhD
Karen L. Bell, MD

Washington University, St. Louis:
Beau Ances, MD
John C. Morris, MD
Maria Carroll, RN, MSN
Mary L. Creech, RN, MSW
Erin Franklin, MS, CCRP
Mark A. Mintun, MD-Past Investigator
Stacy Schneider, APRN, BC, GNP - Past Investigator
Angela Oliver, RN, BSN, MSG-Past Investigator

University of Alabama - Birmingham:
Daniel Marson, JD, PhD
David Geldmacher, MD
Marissa Natelson Love, MD
Randall Griffith, PhD, ABPP-Past Investigator
David Clark, MD-Past Investigator
John Brockington, MD-Past Investigator
Erik Roberson, MD-Past Investigator

Mount Sinai School of Medicine:
Hillel Grossman, MD
Effie Mitsis, PhD

Rush University Medical Center:
Raj C. Shah, MD
Leyla deToledo-Morrell, PhD-Past Investigator

Wien Center:
Ranjan Duara, MD
Maria T. Greig-Custo, MD
Warren Barker, MA, MS

Johns Hopkins University:
Marilyn Albert, PhD
Chiadi Onyike, MD
Daniel D’Agostino II, BS
Stephanie Kielb, BS - Past Investigator

New York University:
Martin Sadowski, MD, PhD
Mohammed O. Sheikh, MD
Anaztasia Ulysse
Mrunalini Gaikwad

Duke University Medical Center:
P. Murali Doraiswamy, MBBS, FRCP
Jeffrey R. Petrella, MD
Salvador Borges-Neto, MD
Terence Z. Wong, MD-Past Investigator
Edward Coleman-Past Investigator

University of Pennsylvania:
Steven E. Arnold, MD
Jason H. Karlawish, MD
David A. Wolk, MD
Christopher M. Clark, MD

University of Kentucky:
Charles D. Smith, MD
Greg Jicha, MD
Peter Hardy, PhD
Partha Sinha, PhD
Elizabeth Oates, MD
Gary Conrad, MD

University of Pittsburgh:
Oscar L. Lopez, MD
MaryAnn Oakley, MA
Donna M. Simpson, CRNP, MPH

University of Rochester Medical Center:
Anton P. Porsteinsson, MD
Bonnie S. Goldstein, MS, NP
Kim Martin, RN
Kelly M. Makino, BS-Past Investigator
M. Saleem Ismail, MD-Past Investigator
Connie Brand, RN-Past Investigator

University of California, Irvine:
Steven G. Potkin, MD
Adrian Preda, MD
Dana Nguyen, PhD

University of Texas Southwestern Medical School:
Kyle Womack, MD
Dana Mathews, MD, PhD
Mary Quiceno, MD

Emory University:
Allan I. Levey, MD, PhD
James J. Lah, MD, PhD
Janet S. Cellar, DNP, PMHCNS-BC

University of Kansas, Medical Center:
Jeffrey M. Burns, MD
Russell H. Swerdlow, MD
William M. Brooks, PhD

University of California, Los Angeles:
Liana Apostolova, MD
Kathleen Tingus, PhD
Ellen Woo, PhD
Daniel H.S. Silverman, MD, PhD
Po H. Lu, PsyD - Past Investigator
George Bartzokis, MD - Past Investigator

Mayo Clinic, Jacksonville:
Neill R Graff-Radford, MBBCH, FRCP (London)
Francine Parfitt, MSH, CCRC
Kim Poki-Walker, BA

Indiana University:
Martin R. Farlow, MD
Ann Marie Hake, MD
Brandy R. Matthews, MD - Past Investigator
Jared R. Brosch, MD Scott Herring, RN, CCRC

Yale University School of Medicine:
Christopher H. van Dyck, MD
Richard E. Carson, PhD
Martha G. MacAvoy, PhD
Pradeep Varma, MD

McGill Univ., Montreal-Jewish General Hospital:
Howard Chertkow, MD
Howard Bergman, MD
Chris Hosein, MEd

Sunnybrook Health Sciences, Ontario:
Sandra Black, MD, FRCPC
Bojana Stefanovic, PhD
Curtis Caldwell, PhD

U.B.C. Clinic for AD & Related Disorders:
Ging-Yuek Robin Hsiung, MD, MHSc, FRCPC
Benita Mudge, BS Vesna Sossi, PhD
Howard Feldman, MD, FRCPC - Past Investigator
Michele Assaly, MA - Past Investigator

Cognitive Neurology - St. Joseph's, Ontario:
Elizabeth Finger, MD
Stephen Pasternack, MD, PhD
Irina Rachisky, MD
Dick Trost, PhD-Past Investigator
Andrew Kertesz, MD-Past Investigator

Cleveland Clinic Lou Ruvo Center for Brain Health:
Charles Bernick, MD, MPH
Donna Munic, PhD

Northwestern University:
Marek-Marsel Mesulam, MD
Emily Rogalski, PhD
Kristine Lipowski, MA
Sandra Weintraub, PhD
Borna Bonakdarpour, MD
Diana Kerwin, MD-Past Investigator
Chuang-Kuo Wu, MD, PhD-Past Investigator
Nancy Johnson, PhD-Past Investigator

Premiere Research Inst (Palm Beach Neurology):
Carl Sadowsky, MD
Teresa Villena, MD

Georgetown University Medical Center:
Raymond Scott Turner, MD, PhD
Kathleen Johnson, NP
Brigid Reynolds, NP

Brigham and Women's Hospital:
Reisa A. Sperling, MD
Keith A. Johnson, MD
Gad Marshall, MD

Stanford University:
Jerome Yesavage, MD
Joy L. Taylor, PhD
Barton Lane, MD
Allyson Rosen, PhD-Past Investigator
Jared Tinklenberg, MD-Past Investigator

Banner Sun Health Research Institute:
Marwan N. Sabbagh, MD
Christine M. Belden, PsyD
Sandra A. Jacobson, MD
Sherye A. Sirrel, CCRC

Boston University:
Neil Kowall, MD
Ronald Killiany, PhD
Andrew E. Budson, MD
Alexander Norbash, MD - Past Investigator
Patricia Lynn Johnson, BA – Past Investigator

Howard University:
Thomas O. Obisesan, MD, MPH
Saba Wolday, MSc
Joanne Allard, PhD

Case Western Reserve University:
Alan Lerner, MD
Paula Ogrocki, PhD
Curtis Tatsuoka, PhD
Parianne Fatica, BA, CCRC

University of California, Davis - Sacramento:
Evan Fletcher, PhD
Pauline Maillard, PhD
John Olichney, MD
Charles DeCarli, MD - Past Investigator
Owen Carmichael, PhD - Past Investigator

Neurological Care of CNY:
Smita Kittur, MD – Past Investigator

Parkwood Hospital:
Michael Borrie, MB
ChB T-Y Lee, PhD
Dr Rob Bartha, PhD

University of Wisconsin:
Sterling Johnson, PhD
Sanjay Asthana, MD
Cynthia M. Carlsson, MD, MS

University of California, Irvine - BIC:
Steven G. Potkin, MD
Adrian Preda, MD
Dana Nguyen, PhD

Banner Alzheimer's Institute:
Pierre Tariot, MD
Anna Burke, MD
Ann Marie Milliken, NMD
Nadira Trncic, MD, PhD, CCRC-Past Investigator
Adam Fleisher, MD-Past Investigator
Stephanie Reeder, BA-Past Investigator

Dent Neurologic Institute:
Vernice Bates, MD
Horacio Capote, MD
Michelle Rainka, PharmD, CCRP

Ohio State University:
Douglas W. Scharre, MD
Maria Kataki, MD, PhD
Brendan Kelley, MD

Albany Medical College:
Earl A. Zimmerman, MD
Dzintra Celmins, MD
Alice D. Brown, FNP

Hartford Hospital, Olin Neuropsychiatry Research Center:
Godfrey D. Pearlson, MD
Karen Blank, MD
Karen Anderson, RN

Dartmouth-Hitchcock Medical Center:
Laura A. Flashman, PhD
Marc Seltzer, MD
Mary L. Hynes, RN, MPH
Robert B. Santulli, MD - Past Investigator

Wake Forest University Health Sciences:
Kaycee M. Sink, MD, MAS
Leslie Gordineer
Jeff D. Williamson, MD, MHS - Past Investigator
Pradeep Garg, PhD - Past Investigator
Franklin Watkins, MD - Past Investigator

Rhode Island Hospital:
Brian R. Ott, MD
Geoffrey Tremont, PhD
Lori A. Daiello, Pharm.D, ScM

Butler Hospital:
Stephen Salloway, MD, MS
Paul Malloy, PhD
Stephen Correia, PhD

UC San Francisco:
Howard J. Rosen, MD
Bruce L. Miller, MD
David Perry, MD

Medical University South Carolina:
Jacobo Mintzer, MD, MBA
Kenneth Spicer, MD, PhD
David Bachman, MD

St. Joseph’s Health Care:
Elizabeth Finger, MD
Stephen Pasternak, MD
Irina Rachinsky, MD
John Rogers, MD
Andrew Kertesz, MD - Past Investigator
Dick Drost, MD - Past Investigator

Nathan Kline Institute
Nunzio Pomara, MD
Raymundo Hernando, MD
Antero Sarrael, MD

University of Iowa College of Medicine
Susan K. Schultz, MD
Karen Ekstam Smith, RN
Hristina Koleva, MD
Ki Won Nam, MD
Hyungsub Shim, MD-Past Investigator

Cornell University
Norman Relkin, MD, PhD
Gloria Chiang, MD
Michael Lin, MD
Lisa Ravdin, PhD

University of South Florida: USF Health Byrd Alzheimer’s Institute
Amanda Smith, MD
Balebail Ashok Raj, MD
Kristin Fargher, MD-Past Investigator

### ADNI Participating Institutions

Johns Hopkins University; Washington University, St. Louis; University of California, Los Angeles; University of Pennsylvania; Cleveland Clinic Lou Ruvo Center for Brain Health; Sunnybrook Health Sciences Centre; Parkwood Hospital; University of California, San Diego; University of Kansas; Dent Neurologic Institute; McGill University / Jewish General Hospital Memory Clinic; Rush University Medical Center; Baylor College of Medicine; Duke University Medical Center; Wein Center for Clinical Research; Indiana University; St. Joseph’s Health Center–Cognitive Neurology; Banner Alzheimer’s Institute; New York University Medical Center; Mayo Clinic, Jacksonville; Mount Sinai School of Medicine; University of Michigan, Ann Arbor; University of British Columbia, Clinic for AD & Related; University of Wisconsin; Oregon Health and Science University; Northwestern University; Boston University; Case Western Reserve University; Emory University; University of Pittsburgh; Brigham and Women’s Hospital; University of Alabama, Birmingham; Medical University of South Carolina; University of California, Irvine; Howard University; University of California, Davis; Rhode Island Hospital; Mayo Clinic, Rochester; Nathan Kline Inst. for Psychiatric Rsch; University of Rochester Medical Center; University of California, Irvine (BIC); The Weill Cornell Memory Disorders Program; Georgetown University; University of California, San Francisco; Banner Sun Health Research Institute; Premiere Research Institute; Butler Hospital Memory and Aging Program; Dartmouth Medical Center; Ohio State University; University of Southern California; University of Iowa; Wake Forest University Health Sciences; University of Kentucky; University of South Florida, Tampa; Columbia University; Yale University School of Medicine; University of Texas, Southwestern MC; Stanford / PAIRE; Albany Medical College.

The list is also available online at http://adni.loni.usc.edu/about/centerscores/study-sites/.

